# Glycolysis Fuels Phosphoinositide 3-Kinase Signaling to Bolster T Cell Immunity

**DOI:** 10.1101/2020.03.12.989707

**Authors:** Ke Xu, Na Yin, Min Peng, Efstathios G. Stamatiades, Amy Shyu, Peng Li, Xian Zhang, Mytrang H. Do, Zhaoquan Wang, Kristelle J. Capistrano, Chun Chou, Andrew G. Levine, Alexander Y. Rudensky, Ming O. Li

## Abstract

Infection triggers clonal expansion and effector differentiation of microbial antigen-specific T cells in association with metabolic reprograming. Here, we show that the glycolytic enzyme lactate dehydrogenase A (LDHA) is induced in CD8^+^ T effector cells via phosphoinositide 3-kinase (PI3K)-dependent mechanisms. In turn, ablation of LDHA inhibits PI3K-dependent phosphorylation of Akt and its transcription factor target Foxo1, causing defective antimicrobial immunity. LDHA deficiency cripples cellular redox control and diminishes glycolytic adenosine triphosphate (ATP) production in effector T cells, resulting in attenuated PI3K signaling. Thus, nutrient metabolism and growth factor signaling are highly integrated processes with glycolytic ATP serving as a rheostat to gauge PI3K/Akt/Foxo1 signaling in T cell immunity control. Such a bioenergetics mechanism of signaling regulation implies a root cause for the century-old phenomenon of Warburg effect, and could guide development of novel therapeutics for infectious diseases and cancer.

**One Sentence Summary:** A PI3K and LDHA circuit enables T cell immunity

## Main Text

In response to infection, microbial antigen-specific naïve T (Tn) cells undergo clonal expansion and effector T (Teff) cell differentiation to eliminate pathogens. Such a T cell fate specification is driven by T cell receptor (TCR) and accessory growth factor receptor signals that reprogram gene expression and nutrient metabolism (*1–4*). Yet, how metabolic processes are adapted to support Teff cell responses remains poorly understood. LDHA, the enzyme that converts pyruvate to lactate and defines the biochemical reaction of aerobic glycolysis, accounts for ∼70% glucose consumed by activated CD4^+^ T cells, and promotes IFN-γ expression through an epigenetic mechanism (*5*). However, glycolysis control of CD8^+^ T cell response has yet to be defined, with a recent study doubting its importance (*6*). To this end, we utilized a model of intracellular bacterial infection with *Listeria monocytogenes* expressing chicken ovalbumin protein (LM-OVA) as an antigen (*7*). H-2K^b^-OVA^+^ Teff cells isolated from day 7-infected mice expressed approximate 5-fold higher LDHA than CD8^+^ Tn cells (Fig. 1A and fig. S1A). Activation of CD8^+^ Tn cells with increasing doses of anti-CD3 progressively induced LDHA expression (Fig. 1B and fig. S1B), in line with dose-dependent induction of the LDHA transcriptional regulator c-Myc (*8*) (Fig. 1B). Notably, while proximal TCR signaling, marked by phospho-Zap70 and phospho-LAT, was saturated at low anti-CD3 doses (fig. S1C), PI3K-dependent Akt phosphorylation showed a dose response similar to c-Myc and LDHA (Fig. 1B). Indeed, blockade of PI3K signaling repressed c-Myc and LDHA expression (Fig.1C and fig. S1D). These observations, together with previous findings that PI3K/Akt signaling promotes plasma membrane localization of the glucose transporter Glut1 (*9*), support a multifaceted role for PI3K in control of glucose metabolism in Teff cells.

**Fig. 1.**
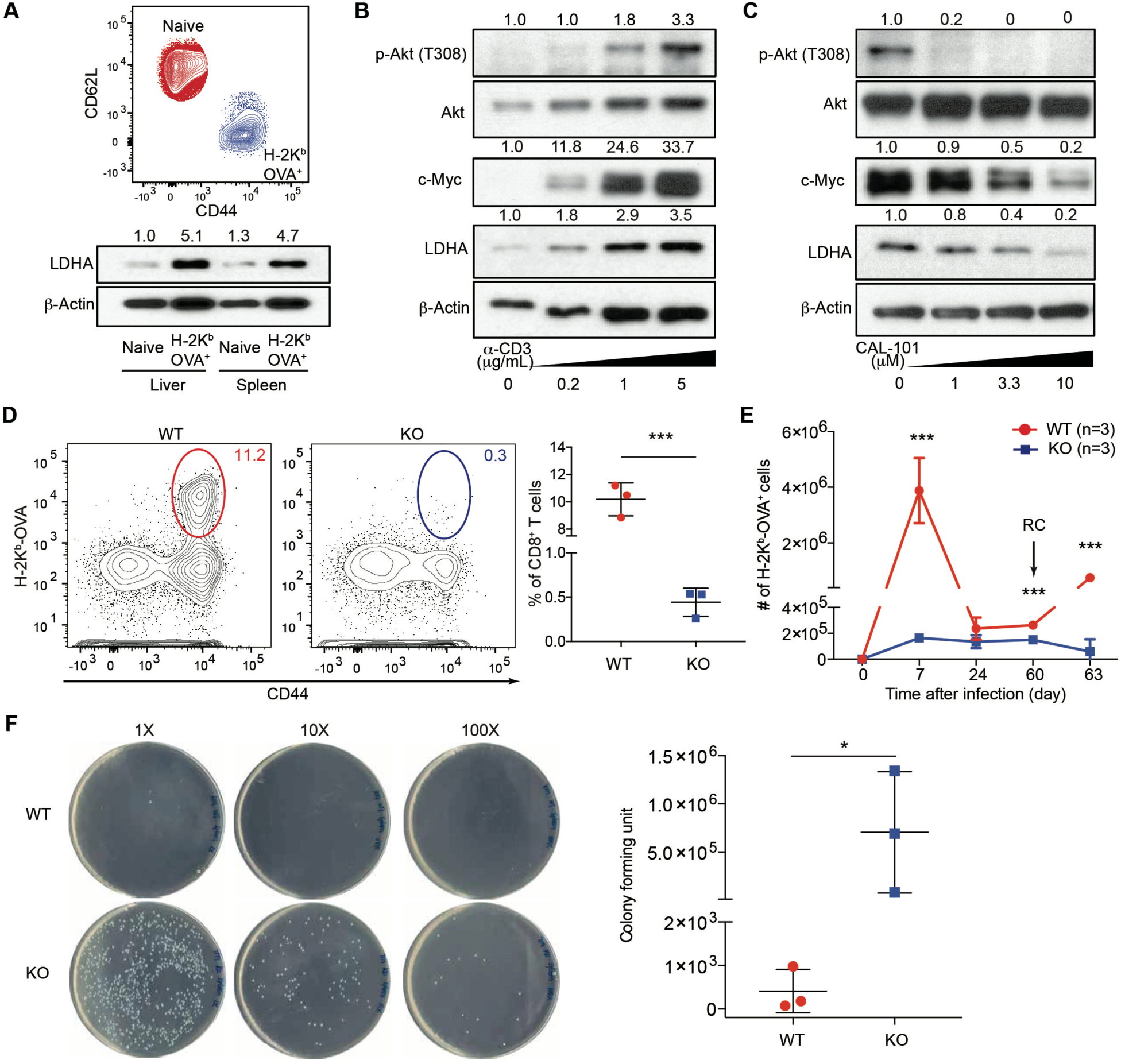
PI3K-dependent expression of LDHA in CD8^+^ effector T cells is essential for anti-bacterial immunity. (**A**), Naïve and H-2K^b^-OVA^+^ CD8^+^ T cells were isolated from spleens and livers of mice infected with *Listeria*-OVA (LM-OVA) for 7 days (n=3). Cellular extracts were subject for immunoblotting. Normalized expression of LDHA to β-Actin was marked. (**B**) Naïve CD8^+^ T cells were stimulated with increasing doses of anti-CD3 in the presence of 2 μg/ml anti-CD28, and 100 U/ml IL-2 for 24 hours. Cellular extracts were immunoblotted for p-Akt (T308), Akt, c-Myc, LDHA, and β-Actin. Normalized expression of p-Akt (T308) to Akt as well as c-Myc and LDHA to β-Actin were marked. (**C**) Naïve CD8^+^ T cells were stimulated with 5 μg/ml anti-CD3, 2 μg/ml anti-CD28, and 100 U/ml IL-2 in the absence or presence of increasing doses of the PI3K inhibitor, CAL-101, for 24 hours. Cellular extracts were immunoblotted for p-Akt (T308), Akt, c-Myc, LDHA, and β-Actin. Normalized expression of p-Akt (T308) to Akt as well as c-Myc and LDHA to b-Actin were marked. (**D** to **F**) *Ldha*^fl/fl^ (wild-type, WT) and Tbx21^Cre^*Ldha*^fl/fl^ (knockout, KO) mice were infected with 5×10^3^ CFU LM-OVA followed or not by re-challenge with 1×10^5^ CFU LM-OVA. (**D**) Representative flow cytometry plots and frequencies of splenic H-2K^b^-OVA^+^ CD8^+^ T cells in WT and KO mice 7 days post LM-OVA infection. (**E**) H-2K^b^-OVA^+^ CD8^+^ T cells from WT and KO mice were enumerated 7, 24, and 60 days post-infection. Day-60-infected mice were also re-challenged (RC) with LM-OVA and splenic H-2K^b^-OVA^+^ CD8^+^ T cells were counted day-3 post-secondary infection (n= 3 per genotype, mean ± SD). (**F**) Splenic bacterial burden from day-3-re-challenged WT and KO mice was assessment by colony forming unit assay (n=3 per group, mean ± SD). Unpaired t tests for the measurements between the two groups (**D** to **F**): *p<0.05; ***p<0.001.

To investigate LDHA function in CD8^+^ Teff cells, we utilized Tbx21^Cre^ mice that specifically targeted H-2K^b^-OVA^+^ Teff cells, revealed by ZsGreen, a Cre recombinase activity reporter (figs. S2A to C). *Ldha*^fl/fl^ (designated as wild-type, WT) and Tbx21^Cre^*Ldha*^fl/fl^ (designated as knock-out, KO) mice were infected with LM-OVA followed by re-challenge, as protective immunity is dependent on CD8^+^ T cells during a recall response (*7*). We found that KO mice had barely detectable antigen-specific Teff cells in contrast to robust expansion in WT mice (Fig. 1D). Defective T cell responses were observed throughout the course of bacterial infection (Fig. 1E), and pathogen was efficiently cleared from the spleens of WT, but not KO, mice (Fig. 1F and fig. S2D). Similarly defective T cell responses were detected in livers of KO mice (figs. S2E to G).

To extend these findings to a viral infection model, WT and KO mice were challenged with lymphocytic choriomeningitis virus (LCMV) clone Armstrong. Diminished expansion of splenic and liver CD8^+^ Teff cells against LCMV GP33 and NP396 antigens were observed in KO mice at day 7 post-infection (figs. S3A to B). While GP33- and NP396-reactive CD8^+^ T cells from WT mice displayed a fully activated phenotype characterized by high CD44 and low CD62L expression, viral antigen-specific CD8^+^ T cells recovered from KO mice expressed high levels of CD62L (figs. S3C to F). LCMV-reactive CD8^+^ Teff cells exhibit phenotypical heterogeneity including a subset with high KLRG1 and low CD127 expression defined as short lived effector cells (SLEC), and a KLRG-1^Lo^CD127^Hi^ subset defined as long-lived memory precursors cells (MPEC) (*10*). LDHA-deficiency resulted in higher proportions of GP33- and NP396-reactive MPECs at the expense of SLECs (figs. S3C to F). Similarly, defective CD62L downregulation and KLRG1 upregulation were observed in H-2K^b^-OVA^+^ Teff cells recovered from KO mice infected with LM-OVA (figs. S4A to B).

To further probe CD8^+^ T cell-intrinsic defects caused by LDHA deficiency, we crossed Tbx21^Cre^*Ldha*^fl/fl^ mice to the OT-I TCR-transgenic background that recognizes OVA SIINFEKL peptide. Congenically marked naïve WT and KO OT-I T cells were mixed at a 1: 1 and co-transferred to receipt mice that were subsequently infected with LM-OVA and monitored for the recovery and phenotype of the transferred T cells (fig. S5A). Time-dependent out-competition of KO OT-I T cells by WT OT-I T cells was observed in the spleen and liver of recipient mice (Fig. 2A and fig. S5B), which was associated with diminished proliferation (Fig. 2B). Attenuated CD62L downregulation and phenotypic skewing towards MPECs were also detected in KO OT-I cells (Fig. 2C and fig. S5C). Together, these observations demonstrate that LDHA promotes expansion and phenotypic diversification of CD8^+^ Teff cells in infection.

**Fig. 2.**
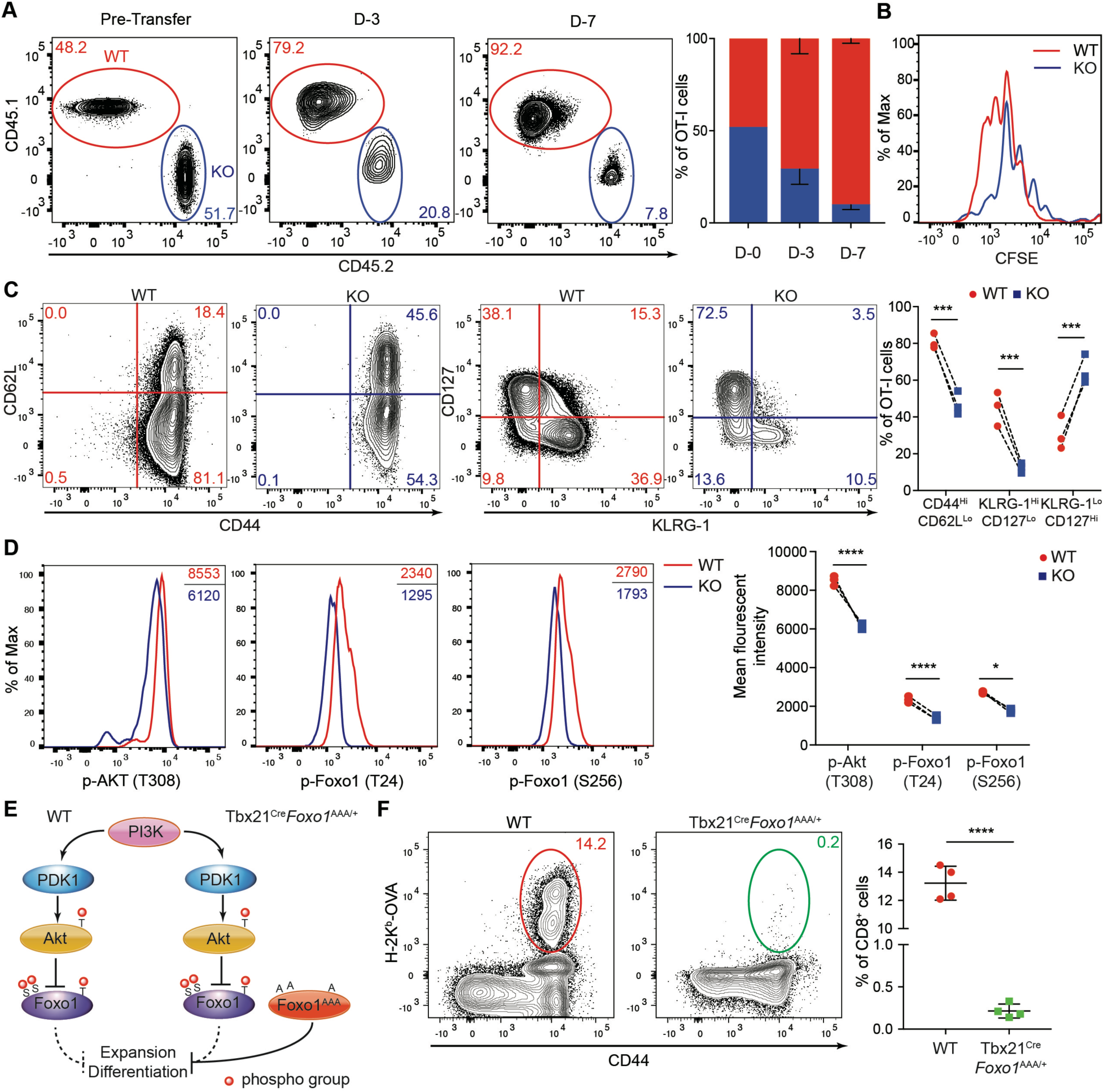
LDHA deficiency impairs Akt and Foxo1 phosphorylation causing defective CD8^+^ T cell expansion and differentiation. **(A** to **D)** Congenically marked naïve OT-I T cells from *Ldha*^fl/fl^ (wild-type, WT) OT-I and Tbx21^Cre^*Ldha*^fl/fl^ (knockout, KO) OT-I mice were CFSE-labeled, mixed at a 1:1 ratio and transferred into WT recipient mice followed by infection with 5×10^3^ CFU LM-OVA. (**A**) Representative plots and ratios (mean ± SD) of splenic WT and KO OT-I T cells from recipient mice day-3 and day-7 post-infection (n=3). (**B**) Proliferation of splenic WT and KO OT-I T cells at day-3 post-infection was assessed by CFSE dilution (n=3). (**C**) Representative flow cytometry plots of CD44, CD62L, CD127, and KLRG-1 expression and the percentages of CD44^Hi^CD62L^Lo^, KLRG-1^Hi^CD127^Lo^, and KLRG-1^Lo^CD127^Hi^ populations among splenic WT and KO OT-I T cells from recipient mice day-7 post-infection (n=3). (**D**) Representative flow cytometry plots and mean florescent intensities of p-Akt (T308), p-Foxo1 (T24), and p-Foxo1 (S256) in splenic WT and KO OT-I T cells from recipient mice day-7 post-infection (n=3). (**E**) A schematic of the PI3K-Akt-Foxo1 signaling pathway. Foxo1^AAA^ was engineered to resist AKT-mediated repression by replacing three phosphorylation sites T24, S256, and S319 with alanine. (**F**) Representative flow cytometry plots and frequencies of splenic H-2K^b^-OVA^+^ CD8^+^ T cells in WT and Tbx21^Cre^*Foxo1*^AAA/+^ mice 7 days post LM-OVA infection (n=4 per genotype, mean ± SD). Paired t tests for the measurements between the two groups (**C** and **D**): *p<0.05; ***p<0.001; ****p<0.0001; unpaired t test for the measurements between the two groups (**F**): ****p<0.0001.

CD8^+^ Teff cell heterogeneity is governed by the strength of TCR and accessory growth factor signaling that modulates the transcriptional programs of T cells (*11–13*). Notably, the transcription factor Foxo1 promotes CD62L expression and KLRG-1^Lo^CD127^Hi^ subset generation (*7, 14, 15*), with its nuclear activity negatively regulated by Akt-induced phosphorylation (*16, 17*). Indeed, diminished phosphorylation of Akt and Foxo1 was observed in KO OT-I T cells (Fig. 2D). Following PI3K activation, Akt and PDK1 are recruited to the plasma membrane whereby PDK1 phosphorylates Akt, and the activated Akt phosphorylates Foxo1 to promote its nuclear exclusion (Fig. 2E). To determine whether compromised Akt-induced Foxo1 phosphorylation might account for Teff cell defects observed in KO mice, we used a mouse strain expressing a mutant form of Foxo1 in which three Akt phosphorylation sites were mutated to alanines (*18*). *Foxo1*^AAA^ expression in CD8^+^ Teff cells was induced by crossing mice carrying the Rosa26-floxed stop *Foxo1*^AAA^ to the Tbx21^Cre^ background (Fig. 2E and fig. S6A). Compared to WT mice, Tbx21^Cre^*Foxo1*^AAA/+^ mice had barely detectable splenic and liver H-2K^b^-OVA^+^ Teff cells that also failed to downregulate CD62L or upregulate KLRG-1 (Fig. 2F and figs. S6B to D). These observations reveal an unexpected function for LDHA in promoting Akt/Foxo1 signaling in CD8^+^ Teff cells with ectopic activation of Foxo1 recapitulating the defects associated with LDHA deficiency.

To determine whether LDHA broadly promoted Akt/Foxo1 signaling in CD8^+^ T cells, we crossed *Ldha*^fl/fl^ mice with CD4^Cre^ transgenic mice to deplete LDHA in Tn cells. WT and KO CD8^+^ Tn cells were acutely stimulated with anti-CD3/CD28, or activated with anti-CD3/CD28 for 3 days followed or not with anti-CD3/CD28 re-stimulation (Fig. 3A). While LDHA deficiency had negligible effects on Akt or Foxo1 phosphorylation in Tn cells (Fig. 3B, lanes 1-4), activated KO T cells had diminished Akt/Foxo1 signaling in the absence or presence of anti-CD3/CD28 re-stimulation (Fig. 3B, lanes 5-8), which was associated with high LDHA expression in activated T cells (Fig. 3B, lanes 1-2 and 5-6). The Akt/Foxo1 signaling defects were specific as proximal TCR signaling, phospho-Zap70 and phospho-LAT, was comparable between WT and KO T cells (Fig. 3B, lanes 13-16).

**Fig. 3.**
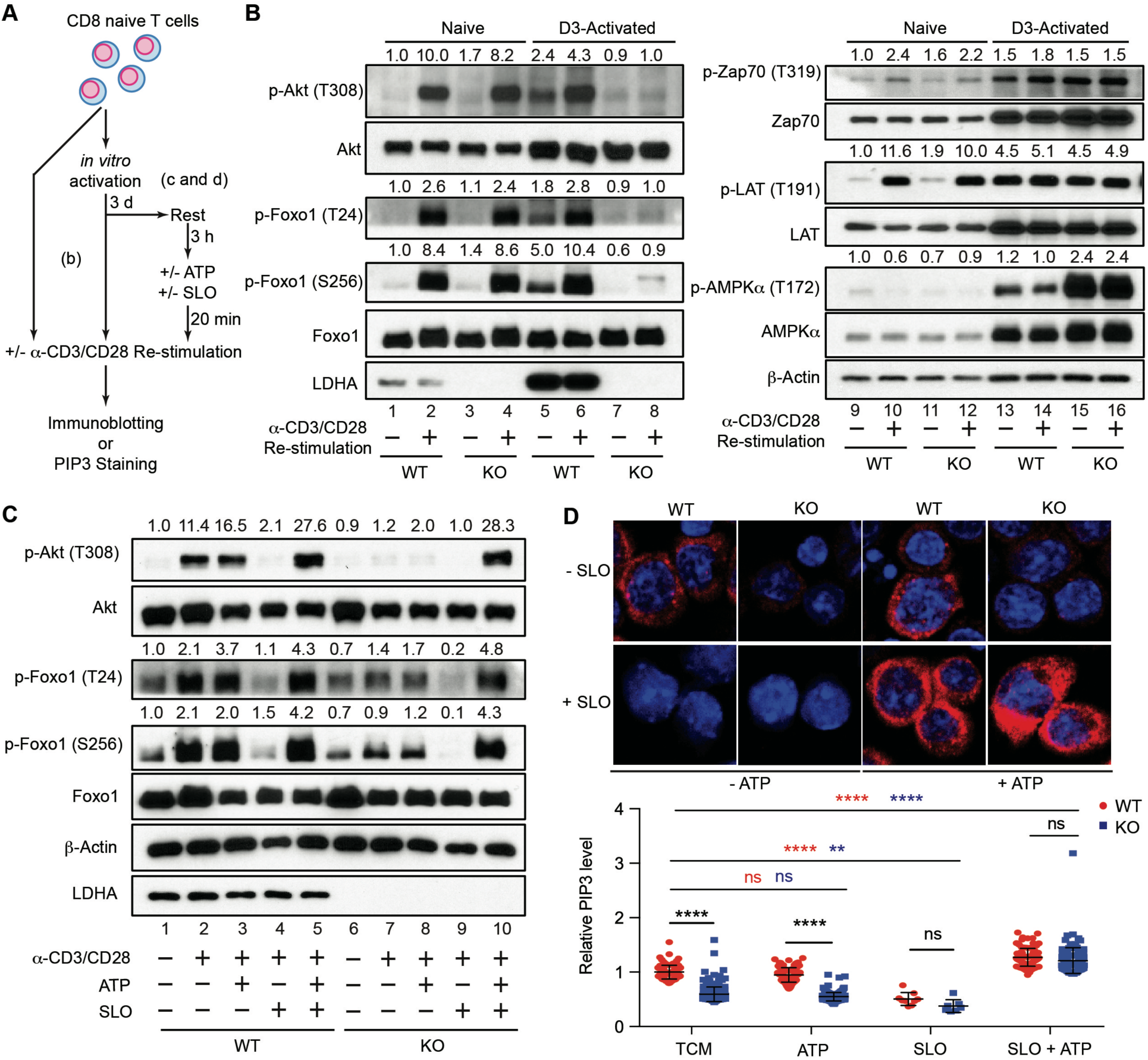
Reduced glycolytic ATP production accounts for defective PI3K/Akt/Foxo1 signaling in LDHA-deficient T cells following antigen stimulation. (**A**) A schematic depicting how naïve CD8^+^ T cells were treated for assessment of kinase signaling and PIP3 generation. (**B**) Naïve CD8^+^ T cells were isolated from *Ldha*^fl/fl^ (wild-type, WT) and CD4^Cre^*Ldha*^fl/fl^ (knockout, KO) mice, and stimulated with 5 μg/ml anti-CD3, 2 μg/ml anti-CD28, and 100 U/ml IL-2 for 3 days. Day-3 activated T cells and freshly isolated naïve WT and KO CD8^+^ T cells were left untreated, or labeled with biotinylated anti-CD3 and anti-CD28 and re-stimulated by streptavidin crosslinking. Cell lysates were prepared and immunoblotted for p-Akt (T308), Akt, p-Foxo1 (T24), p-Foxo1 (S256), Foxo1, LDHA, p-Zap70 (T319), Zap70, p-LAT (T191), LAT, p-AMPKα (T172), AMPKα, and β-Actin. Normalized expression of p-Akt (T308) to Akt, p-Foxo1 (T24) or p-Foxo1 (S256) to Foxo1, p-Zap70 (T319) to Zap70, p-LAT (T191) to LAT, and p-AMPKα (T172) to AMPKα were marked. (**C**) Day-3 activated WT and KO CD8^+^ T cells were collected, rested in RPMI medium for 3 hours, and incubated in the absence or presence of 10 mM ATP and/or 250 U/ml Streptolysin O (SLO). T cells were subsequently re-stimulated with biotinylated anti-CD3 and anti-CD28 through streptavidin crosslinking, and immunoblotted for p-Akt (T308), Akt, p-Foxo1 (T24), p-Foxo1 (S256), Foxo1, LDHA, and β-Actin. Normalized expression of p-Akt (T308) to Akt, p-Foxo1 (T24) or p-Foxo1 (S256) to Foxo1 were marked. (**D**) Representative immunofluorescent images of PIP3 and its quantification in WT and KO CD8^+^ T cells after receiving the indicated treatments as described in (**C**). Unpaired t tests for the measurements between the two groups (**D**): **p<0.01; ****p<0.0001.

LDHA-catalyzed pyruvate to lactate conversion defines an efficient pathway of glucose carbon disposal with its deficiency predicted to diminish glycolysis-associated ATP production. To investigate whether bioenergetics homeostasis was perturbed in LDHA-deficient CD8^+^ T cells, we examined the cellular energy sensor AMPKα that is phosphorylated under energy-stressed conditions (*19*). While AMPKα expression was induced in both WT and KO T cells following activation (Fig. 3B, lanes 13-16), AMPKα phosphorylation was substantially higher in KO T cells (Fig. 3B, lanes 13 and 15). In addition, anti-CD3/CD28 re-stimulation of activated WT T cells led to consistently lower levels of AMPKα phosphorylation (Fig. 3B, lanes 13-14), which was not observed in activated KO T cells (Fig. 3B, lanes 15-16). In agreement with these observations, the ATP level was substantially lower in activated KO T cells than WT T cells (fig. S7A). Collectively, these findings demonstrate a critical function for LDHA in promoting Akt/Foxo1 signaling in Teff cells, which is associated with enhanced ATP production.

To investigate whether reduced ATP production in KO T cells accounted for diminished Akt/Foxo1 signaling, we wished to restore ATP in activated KO T cells. ATP is cell membrane impermeable, and it can be delivered by pore-forming toxins such as Streptolysin-O (SLO) (*20*) (fig. S7B). Day 3-activated CD8^+^ T cells were rested in T cell medium in the absence or presence of ATP and/or SLO, and re-stimulated with anti-CD3/CD28 (Fig. 3A). As expected, anti-CD3/CD28-induced Akt and Foxo1 phosphorylation was strongly inhibited in the absence of LDHA (Fig. 3C, lanes 2 and 7), which was unaltered with the addition of ATP (Fig. 3C, lanes 3 and 8). SLO alone inhibited Akt and Foxo1 phosphorylation (Fig. 3C, lanes 4 and 9), while ATP supplementation together with SLO enhanced Akt/Foxo1 signaling in WT and KO T cells and corrected the defects associated with LDHA deficiency (Fig. 3C, lanes 5 and 10). As the proximal Zap70/LAT-mediated TCR signaling was unaffected in KO T cells (Fig. 3B), we examined whether the differential bioenergetics states impacted the generation of PI3K-catalized phosphatidylinositol (3,4,5)-trisphosphate (PIP3) that serves as a second messenger to recruit PDK1 and Akt to the plasma membrane, a rate-limiting step for PDK1-mediated Akt phosphorylation (*21*). Indeed, KO T cells produced lower amounts of PIP3 which was unaffected by supplementing ATP (Fig. 3D). Inclusion of SLO in the absence or presence of ATP inhibited or enhanced PIP3 generation, respectively, resulting in comparable levels of PIP3 between WT and KO T cells (Fig. 3D). These observations reveal an ATP-sensing property of the PI3K pathway, and suggest that diminished PI3K/Akt/Foxo1 signaling in LDHA-deficient T cells is caused by reduced ATP production.

LDHA-catalyzed pyruvate to lactate conversion regenerates oxidized nicotinamide adenine dinucleotide (NAD^+^) consumed at the glyceraldehyde 3-phosphate dehydrogenase (GAPDH) step of glycolysis (Fig. 4A), which maintains the redox balance and supports high rate of ATP generation via substrate level phosphorylation. Notably, the NAD^+^/NADH ratio was substantially lower in activated KO T cells (Fig. 4B), revealing a critical function for LDHA in maintaining the high oxidation state of T cells. Aside from LDHA-mediated cytosolic pyruvate reduction, the mitochondrion electron transport chain (ETC)-mediated O2 reduction defines an alternative pathway of electron disposal (Fig. 4A). Seahorse extracellular flux analysis showed that compared to day 3-activated WT T cells, KO T cells exhibited a lower level of glucose-induced extracellular acidification rate (ECAR), but a higher baseline oxygen consumption rate (OCR), resulting in a higher OCR/ECAR ratio (Fig. 4C and figs. S8A to B). These findings imply enhanced ETC-mediated redox control in the absence of LDHA.

**Fig. 4.**
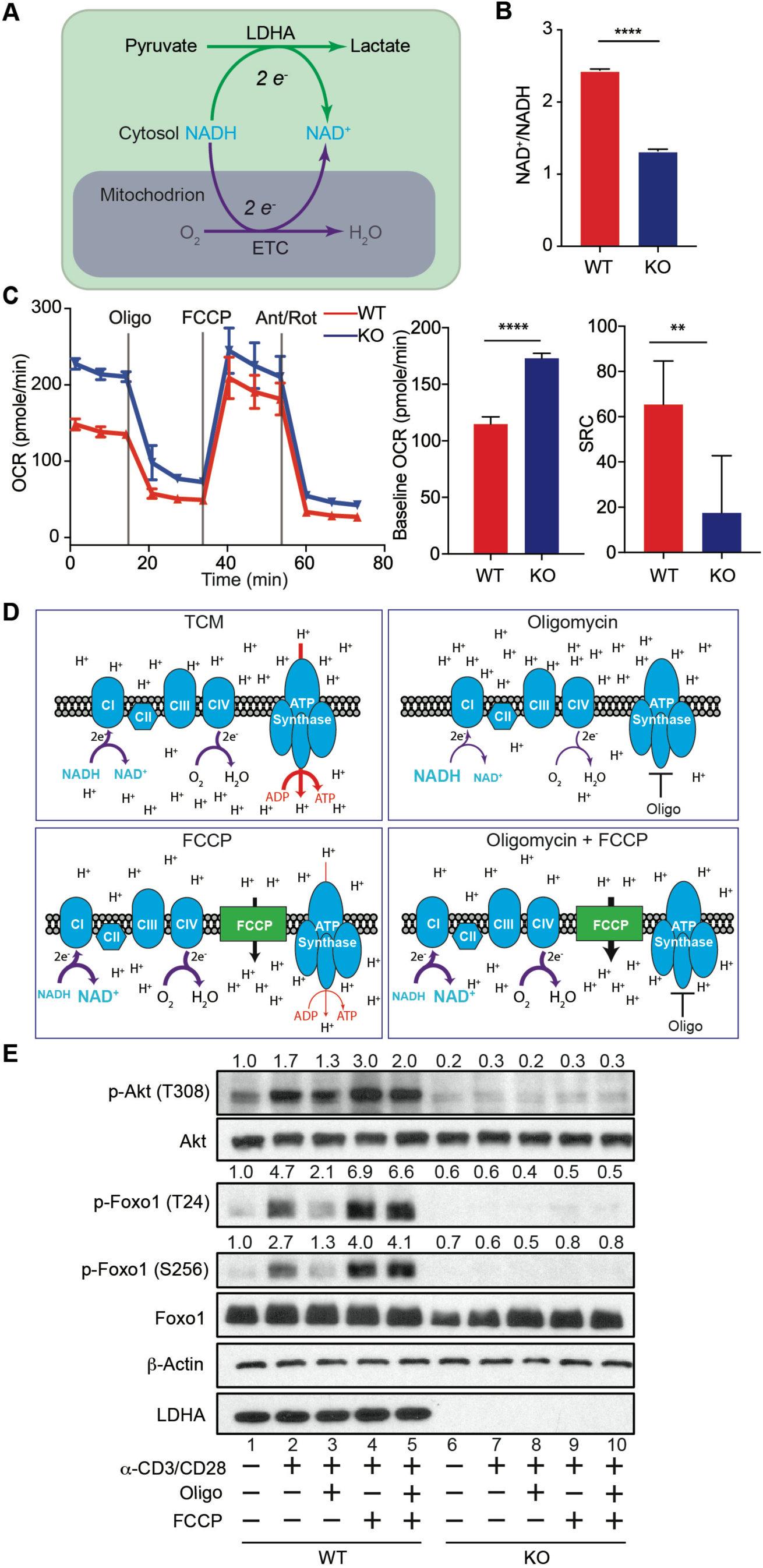
Mitochondrial respiration, but not ATP production, promotes Akt/Foxo1 signaling in activated T cells. (**A**) A schematic depicting electron transfer and disposition by cytosolic LDHA and mitochondrial electron transport chain (ETC) associated with NADH to NAD^+^ conversion. (**B**) Naïve CD8^+^ T cells were isolated from *Ldha*^fl/fl^ (wild-type, WT) and CD4^Cre^*Ldha*^fl/fl^ (knockout, KO) mice, and stimulated with 5 μg/ml anti-CD3, 2 μg/ml anti-CD28, and 100 U/ml IL-2 for 3 days. The amounts of NADH and NAD^+^ were measured. The ratios of NAD^+^ over NADH are plotted. (**C**) Measurements and quantifications of oxygen consumption rate (OCR) and spared respiratory capacity (SRC) of day-3 activated WT and KO CD8^+^ T cells. Sequential chemical treatments are indicated as shown in the graph. Oligo: oligomycin; Ant: antimycin; Rot: rotenone. (n=5 per genotype, mean ± SD) (**D**) A schematic of mitochondrial ETC, proton distribution, and ATP synthetase activity under the indicated treatment conditions. (**E**) Day-3 activated WT and KO CD8^+^ T cells were collected, and incubated in RPMI medium in the absence or presence of oligomycin and/or FCCP. T cells were subsequently re-stimulated with biotinylated anti-CD3 and anti-CD28 through streptavidin crosslinking, and immunoblotted for p-Akt (T308), Akt, p-Foxo1 (T24), p-Foxo1 (S256), Foxo1, LDHA, and β-Actin. Normalized expression of p-Akt (T308) to Akt, p-Foxo1 (T24) or p-Foxo1 (S256) to Foxo1 were marked. Unpaired t tests for the measurements between the two groups (**B** and **C**): **p<0.01; ****p<0.0001.

NADH oxidized by the ETC can be generated through the citric acid cycle, or produced in the cytosol and imported through redox shuttle systems into the mitochondrion (fig. S8C). Both pathways are coupled with generation of a proton gradient for ATP synthesis through oxidative phosphorylation, while the latter can also indirectly support glycolytic ATP production (fig. S8C). To delineate contribution of these two ETC-dependent sources of ATP to Akt/Foxo1 signaling, we utilized oligomycin, an ATP synthase inhibitor that not only blocks mitochondrial ATP synthesis but also suppresses ETC function consequent to mitochondrial hyperpolarization (Fig. 4D and fig. S8C); and FCCP, an mitochondrial proton gradient uncoupler that attenuates mitochondrial ATP synthesis, but enhances ETC activities (Fig. 4D and fig. S8C). As expected, oligomycin diminished OCR in T cells, which was overridden by the addition of FCCP (Fig. 4C). In association with blunted OCR, oligomycin attenuated anti-CD3/CD28-induced Akt and Foxo1 phosphorylation in activated WT T cells (Fig. 4E, lanes 2 and 3), whereas Akt/Foxo1 signaling in KO T cells was barely detectable (Fig. 4E, lanes 7 and 8). Notably, FCCP or oligomycin plus FCCP treatment promoted Akt/Foxo1 signaling in WT, but not KO, T cells (Fig. 4E, lanes 2, 4-5, 7, and 9-10), in agreement with their higher spare respiratory capacity (SRC) (Fig. 4C). Importantly, FCCP-triggered enhancement of Akt and Foxo1 phosphorylation in WT T cells was blocked by ETC complex I and III inhibitors (fig. S8D, lanes 6-10), and the fluctuation of Akt/Foxo1 signaling tracked with OCR as well as ECAR activities, but not with mitochondrial ATP production (Figs. 4C to D and fig. S8E). In contrast to activated T cells, WT and KO CD8^+^ Tn cells exhibited comparable ECAR and OCR activities (figs. S9A to C), and mitochondrial ATP production was essential for anti-CD3/CD28-induced Akt and Foxo1 phosphorylation (fig. S9D). These observations demonstrate that Tn cell to Teff cell transition is associated with bioenergetics reprogramming with Akt/Foxo1 signaling strength modulated selectively by the level of glycolytic ATP, and glycolytic bioenergetics is enabled primarily via LDHA-mediated redox control with ETC playing a subsidiary role.

Antigen stimulation of CD8^+^ T cells induces glucose uptake in association with enhanced effector function (*8, 22–25*); yet, how glucose is metabolized to support Teff cell responses *in vivo* is not well defined. A recent study of glucose tracing in CD8^+^ Teff cells suggested that glucose was primarily used for anabolic processes instead of catabolized through glycolysis (*6*), although the lengthy protocol of cell isolation could underestimate the recovery of lactate that is readily excreted from T cells. Herein, using a genetic model of LDHA deficiency, we found that glycolysis is indispensable for CD8^+^ Teff cell responses to bacterial and viral infections. An important function of LDHA is to augment glycolysis-associated ATP production to fuel PI3K signaling that in turn promotes LDHA expression, constituting a positive feedback loop to reinforce T cell immunity. Notably, glycolytic ATP preferentially supports PI3K signaling in Teff cells, while ATP generated through oxidative phosphorylation fuels PI3K signaling in CD8^+^ Tn cells. Such differential bioenergetics requirements can be consequent to the remodeling of the mitochondrion network in Teff cells (*26*), which may spare it as an energy house for glucose anabolism including generation of acetyl-CoA in support of cytokine expression (*5*). Diminished glycolytic ATP production in LDHA-deficient Teff cells impairs PI3K, but not proximal TCR signaling. Of note, unlike Zap70 with an ATP Km around 3 μM (*27*), PI3K has an ATP Km as high as hundreds μM (*28*), raising the possibility that PI3K is a novel ATP sensor. As enhanced PI3K signaling is a major oncogenic event (*29*), future studies will uncover whether the positive feedback circuit defined in our study provides a mechanistic explanation for the classical phenomenon of Warburg effect in cancer (*30*). Such a bioenergetics mechanism of signaling regulation may as well guide the development of effective vaccines for infectious diseases and therapeutics for cancer.

## Acknowledgments

The authors thank members of the Li laboratory and Morgan Huse for helpful discussions.

## Author Contributions

M.O.L. conceived the project. K.X. designed and performed most experiments with input from M.O.L., N.Y., and M.P. K.X. and E.G.S. conducted the IF experiments. K.X. and A.S. performed the FACS sorting experiments. P.L., X.Z., M.H.D., Z.W., and C.C. assisted with the design of experiments. K.J.C. helped managing the mouse colony. and A.G.L. and A.Y.R. provided critical mouse strains. K.X. wrote and M.O.L. edited the manuscript.

## Data and materials availability

All data is available in the main text or the supplementary materials. All materials used in the research are available upon request.

## Supplementary Materials

### Materials

#### Mice

B6 CD45.1, B6 CD4^Cre^, B6 OT-I and *ZsGreen* reporter mice were purchased from Jackson Laboratory. *Ldha*^fl/fl^ and *Foxo1*^AAA/+^mouse was generated in the laboratory of Ming O. Li (*1, 2*) and B6 Tbx21^tdTomato-T2A-cre^ was generated in the laboratory of Alexander Y. Rudensky (*3*). All *in vivo* mouse experimental procedures were performed under Sloan Kettering Institute (SKI) Institutional Animal Care and Utilization Committee (IACUC) - approved protocols including: *Listeria monocytogenes*-OVA and LCMV (Clone Armstrong) infection are conformed to all relevant regulatory standards. Mice were housed in designated specific pathogen-free animal facilities regarding to normal and infected condition in ventilated cages with at most 5 animals per cage and provided food and water. In all experiments comparing wild-type to LDHA-deficient, healthy, sex-matched and age-matched mice were used (male and female, 8 to 14 weeks of age unless otherwise stated). No sex differences in T cell phenotype were noted.

#### Antibodies

Mouse flow cytometry antibodies: Antibodies against CD127-AF647 (Clone A7R34; Cat. # 135019), CD45.1-BV421 (Clone A20; Cat. # 110731), CD45.1-BV650 (Clone A20; Cat. # 110735), KLRG-1-BV786 (Clone 2F1; Cat. # 138429), TCRβ-APC-Cy7 (Clone H57-597; Cat. # 109220) were purchased from Biolegend. Antibodies against CD4-BV510 (Clone RM4-4; Cat. # 740105) and CD4-BV786 (Clone GK1.5; Cat. # 563331) were purchased from BD Biosciences. Antibodies against CD44-PerCP-Cy5.5 (Clone IM7; Cat. # 65-0441-U100), CD45.2-APC (Clone 104; Cat. # 20-0454-U100), CD62L-PE-Cy7 (Clone MEL-14; Cat. # 60-0621-U100), CD8α-vF450 (Clone 53-6.7; Cat. # 75-0081-U100), CD8α-PE-Cy7 (Clone 53-6.7; Cat. # 60-0081-U100), CD8α-PerCP-Cy5.5 (Clone 53-6.7; Cat. # 65-0081-U100), TCRβ-FITC (Clone H57-597; Cat. # 35-5961-U100) were purchased from Tonbo. Antibodies against CD8α-Pacific Blue (Clone KT15; Cat. # MCA609GT) was purchased from Bio Rad. Fluorescent-dye-labeled tetramers H-2D^b^-NP396 Alexa 647, H-2D^b^-GP33 BV421 and H-2K^b^-OVA APC were obtained from NIH. H-2K^b^-OVA PE was manufactured by MSKCC Tetramer Core.

Antibodies for immunoblotting and immunofluorescence staining: Monoclonal rabbit anti-pT308 Akt (Clone D25E6; Cat. # 13038S), polyclonal rabbit anti-pS256 Foxo1 (Cat. # 9461S), polyclonal rabbit anti-pT24 Foxo1/pT32 Foxo3a (Cat. # 9464S), polyclonal rabbit anti-AMPKα (Cat. # 2532S), monoclonal rabbit anti-pT172 AMPKα (Clone 40H9; Cat. # 2535S), monoclonal rabbit anti-PAN Akt (Clone C67E7; Cat. # 4691L), monoclonal rabbit anti-Foxo1 (Clone C29H4; Cat. # 2880S), polyclonal rabbit anti-LDHA (Cat. # 2012S), monoclonal mouse anti-β Actin (Clone 8H10D10; Cat. # 3700S), polyclonal rabbit anti-LAT (Cat. # 9166S), polyclonal rabbit anti-pT191 LAT (Cat. # 3584S), polyclonal rabbit anti-Zap70 (Cat. # 2702) and polyclonal rabbit anti-pT319 Zap70/ pT352 Syk (Cat. # 2701S) were purchased from Cell Signaling Technology. Monoclonal anti-Phosphatidylinositol 3,4,5-trisphosphate (Clone RC6F8; Cat. # A-21328) and goat anti-rabbit IgG (H+L) highly cross-adsorbed secondary antibody, Alexa Fluor Plus 647 (Cat. # A32733) were purchased from Invitrogen.

Antibodies for *in vitro* T cell culture: Monoclonal anti-mouse CD28-biotin (Clone 37.51; Cat. # 13-0281-81), monoclonal anti-mouse and CD3ε-biotin (Clone 145-2C11; Functional Grade; Cat. # 36-0031-85) were purchased from Invitrogen. Monoclonal anti-mouse CD28 (Clone 37.51; Cat. # BE0015-1) and monoclonal anti-mouse CD3ε (Clone 145-2C11; Cat. # BE0001-1) were purchased from BioXCell.

### Methods

#### *Listeria Monocytogenes*-OVA (LM-OVA) Infection and OT-I Co-transfer

Frozen aliquot/vial of LM-OVA (stored at −80 °C) was thawed and 200 μl of bacteria was added into 5 ml sterile brain heart infusion (BHI) media. Bacterial culture was incubated 3.5 to 4 hours in 37 °C shaking at 225 rpm. Bacterial OD600 was monitored and ready when reached between 0.1 ∼ 0.2 (OD600 = 0.1 is equal to 1×10^8^ bacteria/ml). Tbx21^Cre^*Ldha*^fl/fl^ and *Ldha*^fl/fl^ mice were intravenously injected with 5 x 10^3^ colony-forming units (CFU) of LM-OVA. Acute antigen-specific CD8^+^ T cell response was assessed on day-7 and memory T cell response was assessed on day-60. Memory T cell recall response was assessed by re-challenging mice with 1×10^5^ CFU LM-OVA 60 days after primary infection. The re-call response was determined 3 days after secondary infection. For OT-I co-transfer experiment, naïve OT-I CD8^+^ T cells were isolated from Tbx21^Cre^*Ldha*^fl/fl^ (knockout, KO) OT-I (CD45.2) and wild-type (WT) OT-I (CD45.1) mice and were labeled with CFSE. WT and KO OT-I cells were counted and mixed 1:1 ratio. WT/KO cell ratio were checked by flow cytometry to make sure the ratio is close to 1:1. Cell mixture was re-counted and diluted to 7.5×10^5^ cells/ml and 200 μl (1.5×10^5^ cells) were injected to WT host mice (CD45.1/CD45.2). On day-1 post cell transfer, mice were infected with 1×10^5^ CFU LM-OVA. Activation, proliferation and differentiation of the transferred T cells were assessed on day-3 and day-7 post infection.

#### Lymphocytic Choriomeningitis Virus (LCMV Clone Armstrong) Infection

*Ldha*^fl/fl^ and Tbx21^Cre^*Ldha*^fl/fl^ mice were infected with LCMV clone Armstrong (1 × 10^5^ PFU) by intra-peritoneal injection to cause acute viral infection. The numbers of antigen-specific CD8^+^ T cells in spleen and liver were determined by MHC tetramer staining on day-7 post infection against the following epitopes: GP33 and NP396.

#### Cell Sorting

Related to Fig. 1A: spleens and livers were isolated from WT and KO mice on day-7 post LM-OVA infection. After gating on DAPI^-^, morphology and singlets, CD45^+^TCRβ^+^CD8^+^ cells were gated as follow: CD44^Hi^H-2K^b^-OVA^+^ (antigen-specific CD8^+^ T cell) and CD44^Lo^H-2K^b^-OVA^-^. From these two parental populations, CD44^Hi^CD62L^Lo^ (activated) and CD44^Lo^CD62L^Hi^ (naïve) T cells were gated respectively and sorted for immunoblotting.

#### Colony Forming Unit (CFU) Assay

3 days after secondary infection of LM-OVA, spleens and livers were taken and a small piece was separated. The total and small piece from each sample were weighted. Small pieces of samples were made into 3 ml of single cell suspension supplemented with 0.1% Triton X-100. Cell suspensions were diluted in sterile PBS and made 10X and 100X suspension. 100 μl of each sample were added to BHI plate and incubated 37 °C overnight. CFU were counted from all plates and calculated back to total CFU/Spleen or CFU/Liver.

#### Flow Cytometry

Tissues from spleens and livers were grinded and filtered through 100 μm mesh to make single cell suspension. Then cells were depleted of erythrocytes by hypotonic lysis. Cells were incubated with specific antibody cocktails for 15 minutes at 4 °C in the presence of anti-FcγR to block FcγR binding. Dead cells were excluded by DAPI (Invitrogen) staining. For tetramer staining, PE-/APC-/BV421 conjugated H-2K^b^-OVA, H-2D^b^-GP33 and H-2D^b^-NP396 tetramers (NIH) was used and incubated for 30 minutes at 4 °C. For phospho-signal flow cytometry, spleens were isolated and immediately meshed into 4% PFA solution for 10 minutes at room-temperature and followed by 1 hour permeabilization with fix/perm buffer at 4 °C. Cells were then washed and incubated with primary phospho- and secondary-antibodies for flow cytometry assay. All samples were acquired with an LSR II flow cytometer (Becton, Dickinson) and analyzed with FlowJo software.

#### *in vitro* T Cell Activation

T cell *in vitro* experiments were performed by using primary mouse T cells isolated from lymph nodes and spleens of male and female mice cultured at 37 °C with 5% CO2 in T cell medium (TCM): RPMI1640 media (Media Preparation Core, SKI) supplemented with 1000X β-mercaptoethanol (Invitrogen), 100X Penicillin/Streptomycin (Gimini Bio), 100X glutamine-MAX (Thermo Fisher) and 10% heat inactivated FBS (Sigma), unless otherwise stated. Naïve (CD62L^+^ CD44^-^) CD8^+^ T cells were isolated from spleens and lymph nodes of *Ldha*^fl/fl^ or CD4^Cre^*Ldha*^fl/fl^ mice with mouse naïve CD8^+^ T cells isolation kits (Stem Cell). Purity was validated by flow cytometry and was over 95%. For regular *in vitro* T cell activation, 0.2-0.3×10^6^ naïve CD8^+^ T cells were cultured in a 24-well plate pre-coated with 5 μg/ml anti-CD3, and 2 μg/ml soluble anti-CD28 and 100 U/ml rhIL-2 (NIH) in TCM unless specific conditions were mentioned. Related to Fig. 1B and fig. S1C: naïve CD8^+^ T cells were cultured with 0, 0.2, 1 or 5 μg/ml anti-CD3, and 2 μg/ml anti-CD28 and 100 U/ml rhIL-2 in TCM for 24 hours. Cells were collected and used to assess protein induction by immunoblotting. Related to Fig. 1C and fig. S1D: naïve CD8^+^ T cells were cultured with 5 μg/ml anti-CD3, 2 μg/ml anti-CD28 and 100 U/ml rhIL-2 in TCM in the presence of 0, 1, 3.3 or 10 μM of PI3K inhibitor, CAL-101 (Cayman Chemical) for 24 hours. Cells were collected and used to assess protein expression by immunoblotting.

#### ATP, NAD^+^ and NADH Measurements

T cells were harvested 3 days post *in vitro* activation, washed with RPMI1640 (without any supplementations) 3 times and counted. Cells were divided into different treatment groups with 1 million cells in each group and rested in 96-well plate for 2 hours at 37 °C. Cells were quickly spun down and lysed in a native lysis buffer (Cell Signaling Technology) for 10 minutes on ice. Cells were centrifuged at 14000 RPM for 10 minutes and the supernatants were collected. ATP concentration was measured according to an user protocol provided by Invitrogen. For NAD^+^/NADH measurements, cells were lysed in a base lysis solution (100 mM sodium carbonate, 20 mM sodium bicarbonate, 10 mM nicotinamide, 0.05% Triton X-100 and 1% DTAB) for 20 minutes on ice. Cells were centrifuged at 14000 RPM for 10 minutes and the supernatants were collected. NAD^+^ and NADH concentrations were measured accordingly to an user protocol provided by Promega.

#### Extracellular Flux Analyses (Seahorse)

Experiments were performed on Agilent Seahorse XF96 bioanalyzer. Freshly isolated CD8^+^ T cells were adhered onto XF96 microplates (1.5×10^5^ cells/well) that had been pre-coated with Cell-Tak adhesive. The plates were quickly centrifuged to immobilize cells. Cells were rested in a non-buffered assay medium in a non-CO_2_ incubator for 30 minutes before the assay. Oxidative phosphorylation associated parameters were determined by Seahorse XF Cell Mito Stress test kit with three injections: 1) 1 μM oligomycin; 2) 0.25 μM FCCP and 3) 0.5 μM rotenone/antimycin A. Glycolysis associated parameters were determined by Seahorse XF Glycolysis Stress test kit with three injections: 1) 10 mM glucose; 2) 1μM oligomycin; 3) 50 mM 2-DG.

#### Acute Anti-CD3/CD28 Cross-linking and ATP Supplementation

In cell re-stimulation experiments, cells were divided into different treatment groups with 1 million cells in each group. After indicated treatments, cells were labeled with biotinylated anti-CD3 and anti-CD28 for 30 minutes at 4 °C. Then, cells were pre-warmed for 2 minutes in a 37 °C water bath and acute streptavidin-biotin cross-linking was performed for additional 5 minutes. Cell lysates were prepared, and subject to immunoblotting. Related to Fig. 3B: naïve and day 3-activated CD8^+^ T cells were collected and washed with RPMI1640 (without any supplementations) 3 times and counted. Cells were re-stimulated and immunoblotting was used for signaling assessment. Related to Fig. 3C: a Streptolysin-O (SLO) based pore-forming ATP delivery system was used as described in previous publications (*4*). Day 3-activated WT and KO CD8^+^ T cells were collected and rested in 96-well plate for 3 hours at 37 °C. Cells were then washed in a Mg^2+^-, Ca^2+^- and ATP-free HBSS buffer twice and incubated for 20 minutes under the following conditions: 1) HBSS; 2) HBSS + ATP (10 mM); 3) HBSS + SLO (250 U/ml); and 4) HBSS + SLO + ATP. Followed the treatments, cells were re-stimulated and immunoblotted for signaling components. Related to Fig. 4E: day 3-activated T cells were harvested and washed with RPMI 3 times and rested in RPMI for 3 hours and followed by further treatments: 1) TCM; 2) TCM + 2 μM oligomycin; 3) TCM + 3 μM FCCP; 4) TCM + 2 μM oligomycin + 3 μM FCCP. Cell signaling were assessed after re-stimulation. Related to fig. S8D: day 3-activated WT T cells were harvested and washed with RPMI 3 times and rested in RPMI for 3 hours and followed by further treatments: 1) TCM; 2) TCM + 2 μM oligomycin; 3) TCM + 3 μM FCCP; 4) TCM + 2 μM oligomycin + 3 μM FCCP with or without Rotenon/Antimycin A. Cell signaling were assessed after re-stimulation. Related to fig. S9D: naïve T cells were stimulated as described for Fig. 4E.

#### Immunofluorescence Staining

Day 3-activated T cells were harvested from 24-well plates, washed with RPMI1640 (without any supplementations) 3 times and counted. Cells were divided into different treatment groups with 1 million cells in each group and rested in 96-well plate for 3 hours at 37 °C. Cells were subsequently washed in an Mg^2+^-, Ca^2+^- and ATP-free HBSS buffer twice and incubated for 20 minutes under the following conditions: 1) HBSS; 2) HBSS + ATP (10 mM); 3) HBSS + SLO (250 U/ml); and 4) HBSS + SLO + ATP. T cells were then incubated with biotinylated anti-CD3 and anti-CD28 for 30 minutes at 4 °C. T cells were transferred in to 8-well Lab-Tek II chamber slide and pre-warmed for 2 minutes in a 37 °C water bath. Acute streptavidin-biotin crosslinking was performed by adding streptavidin for additional 5 minutes and cells were spun down to the bottom. Cells were then fixed with 4% PFA for 10 minutes at room-temperature and washed once with PBS. Cells were subsequently permeabilized with a Fix/Perm buffer (Tonbo) for 1 hour at 4 °C and washed once with PBS. Primary antibody against PIP3 was diluted in the flow cytometry perm buffer (Tonbo) and added to the fixed samples. The primary antibody incubation took place for overnight at 4 °C. Next day, primary antibody was aspirated, and samples were washed twice with PBS. Secondary antibody was then added and incubated for 1 hour at room-temperature. After secondary antibody staining, cells were washed with PBS twice. Samples were incubated with hoechst diluted in PBS for 10 minutes at room-temperature and then washed with PBS twice. Chamber wall was removed after wash. Antifade mountant was added on top of samples and cover slides were applied. The images were taken with the ZEISS LSM 880 microscope under 20X objective magnification.

#### PIP3 Quantification by ImageJ

The .czi files were opened in ImageJ/FIJI (NIH, USA) using the Bio-formats Importer plugin, and the channels were split into separate 8-bit images. Then, a median filter, thresholding, and marker-controlled watershedding were used to create a mask of the segmented nuclei in the DAPI channel. Next, the Analyze Particles algorithm was run on the mask to create regions around each nucleus. These regions were overlaid onto the red channel with thresholding in order to quantify the amount of positive signal per cell.

#### Statistical Analysis

Statistical tests were performed with paired, unpaired t tests or two-way Anova multiple comparison tests. A value of p<0.05 was considered statistically significant. All error bars represent mean ± standard deviation (SD).

**Fig. S1.**
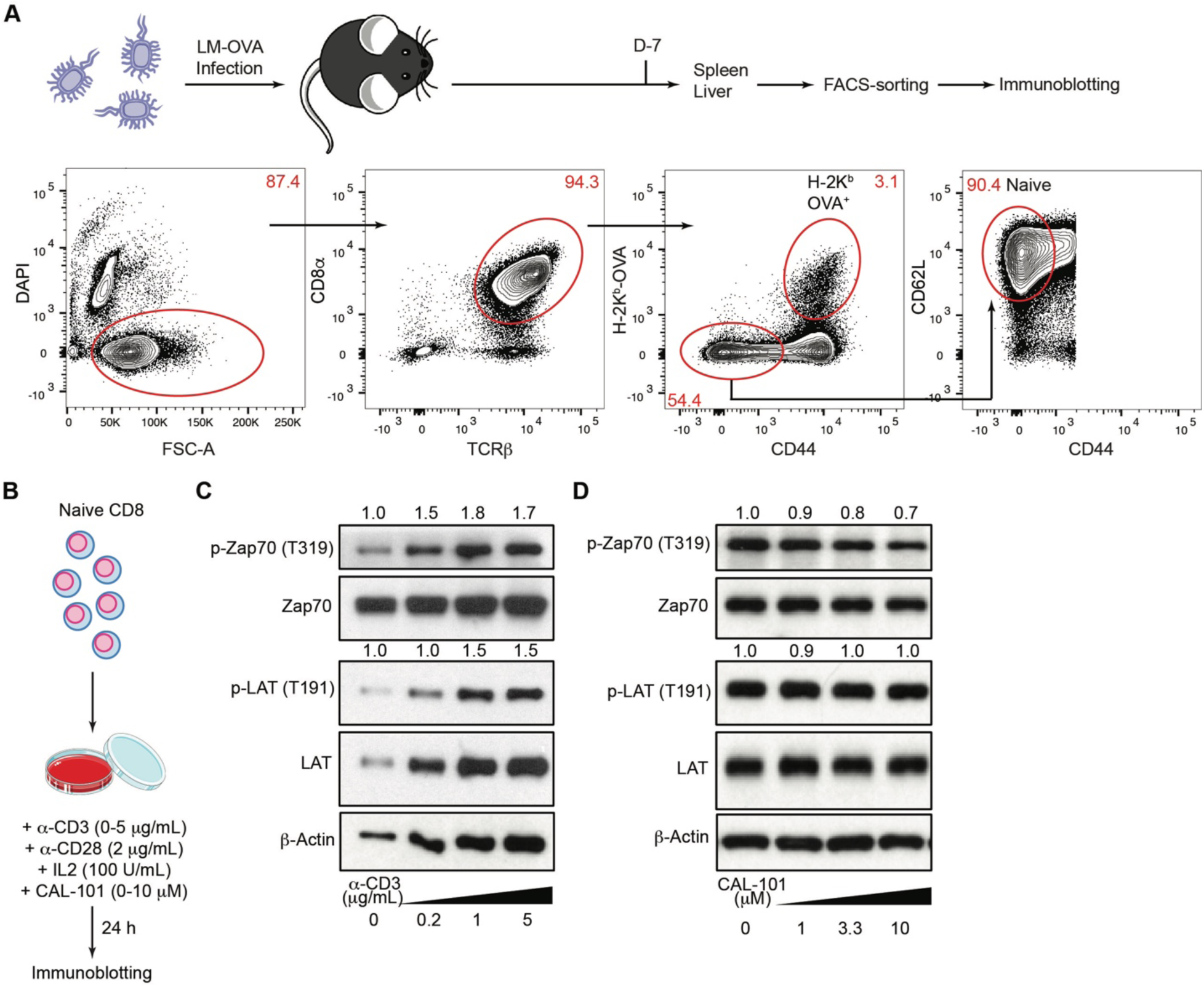
Schematics of cell sorting and cell culture, and characterization of anti-CD3-induced TCR proximal signaling in the absence or presence of PI3K inhibitor. (**A**) A schematic of flow cytometry sorting strategy for naïve and H-2K^b^-OVA^+^ CD8^+^ T cells from wild-type mice infected with LM-OVA for 7 days. (**B**) A schematic of *in vitro* CD8^+^ T cell culture with various doses of anti-CD3 and the PI3K inhibitor, CAL-101 in the presence of 2 μg/ml anti-CD28, and 100 U/ml IL-2. (**C**) Naïve CD8^+^ T cells were stimulated with increasing doses of anti-CD3 in the presence of 2 μg/ml anti-CD28, and 100 U/ml IL-2 for 24 hours. Cellular extracts were immunoblotted for p-Zap70 (T319), Zap70, p-LAT (T191), LAT, and β-Actin. Normalized expression of p-Zap70 (T319) to Zap70 and p-LAT (T191) to LAT were marked. (**D**) Naïve CD8^+^ T cells were stimulated with 5 μg/ml anti-CD3, 2 μg/ml anti-CD28, and 100 U/ml IL-2 in the absence or presence of increasing doses of the PI3K inhibitor, CAL-101, for 24 hours. Cellular extracts were immunoblotted for p-Zap70 (T319), Zap70, p-LAT (T191), LAT, and β-Actin. Normalized expression of p-Zap70 (T319) to Zap70 and p-LAT (T191) to LAT were marked.

**Fig. S2.**
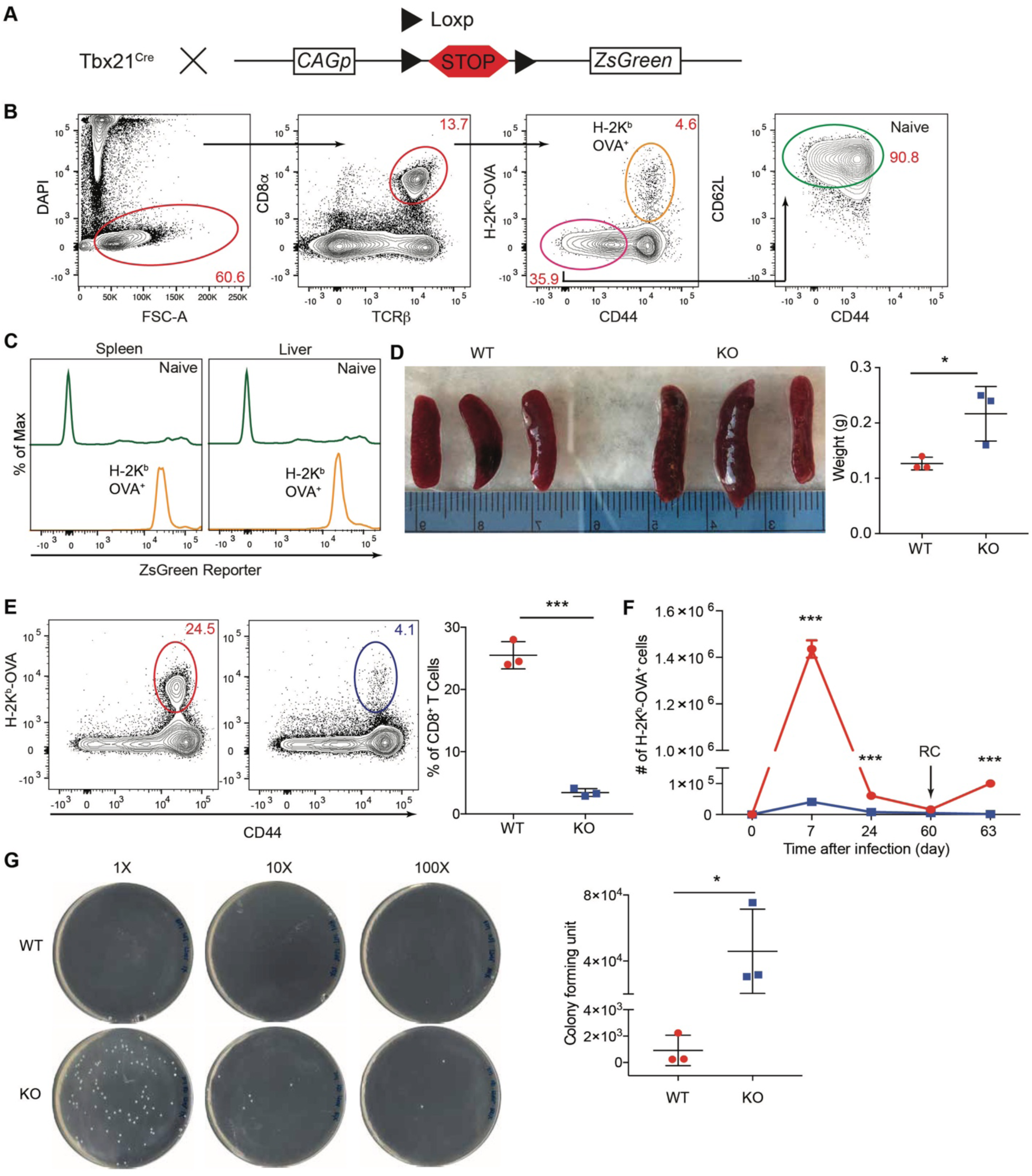
Specificity of the Tbx21^Cre^ line and its use in assessing LDHA control of CD8^+^ effector T cell-mediated anti-bacterial immunity. (**A**) A schematic depicting the mouse breeding strategy and the genomic structure of *ZsGreen* reporter mouse strain. (**B**) Representative flow cytometry gating strategy for naïve and H-2K^b^-OVA^+^ CD8^+^ T cells isolated from the spleen and liver of a Tbx21^Cre^*ZsGreen* reporter mouse 7 days after LM-OVA infection. (**C**) Representative flow cytometry plots of ZsGreen reporter expression in naïve and H-2K^b^-OVA^+^ CD8^+^ T cells isolated from the spleen and liver of a Tbx21^Cre^*ZsGreen* mouse 7 days after LM-OVA infection. (**D**) Photographic illustration and quantification of the weight of spleens isolated from *Ldha*^fl/fl^ (wild-type, WT) and Tbx21^Cre^*Ldha*^fl/fl^ (knockout, KO) mice on day 7 post LM-OVA infection. (**E**) Representative flow cytometry plots and frequencies of liver H-2K^b^-OVA^+^ CD8^+^ T cells in WT and KO mice 7 days post LM-OVA infection (n= 3 per genotype, mean ± SD). (**F**) H-2K^b^-OVA^+^ CD8^+^ T cells from WT and KO mice were enumerated 7, 24, and 60 days post infection. Day-60-infected mice were also re-challenged (RC) with LM-OVA and liver H-2K^b^-OVA^+^ CD8^+^ T cells were counted day-3 post-secondary infection (n= 3 per genotype, mean ± SD). (**G**) Liver bacterial burden from day-3-re-challenged WT and KO mice was assessment by colony forming unit assay (n=3 per group, mean ± SD). Unpaired t tests for the measurements between the two groups (**D** to **G**): *p<0.05; ***p<0.001.

**Fig. S3.**
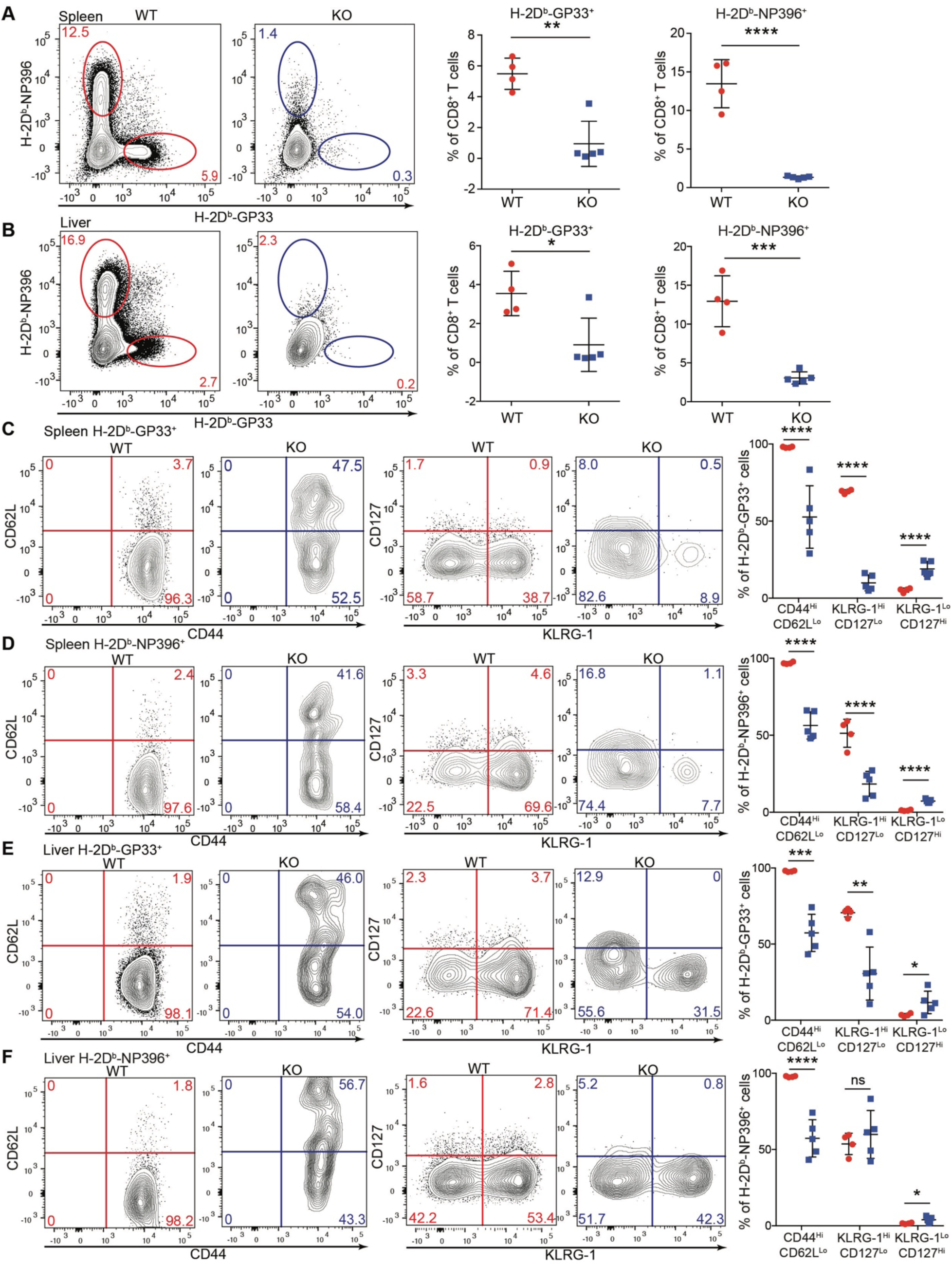
LDHA deficiency impairs CD8^+^ effector T cell response to LCMV infection. *Ldha*^fl/fl^ (wild-type, WT) and Tbx21^Cre^*Ldha*^fl/fl^ (knockout, KO) mice were infected with LCMV Armstrong (1 × 10^5^ PFU) by intra-peritoneal injection. (**A** and **B**) Representative flow cytometry plots and frequencies of H-2D^b^-NP396^+^ and H-2D^b^-GP33^+^ CD8^+^ T cells isolated from the spleens (**A**) and livers (**B**) of WT (n=4) and KO (n=5) mice on day 7 post infection (mean ± SD). (**C** to **F**) Representative flow cytometry plots of CD44, CD62L, CD127, and KLRG-1 expression and the percentages of CD44^Hi^CD62L^Lo^, KLRG-1^Hi^CD127^Lo^, and KLRG-1^Lo^CD127^Hi^ populations among H-2D^b^-GP33^+^ (**C**) and H-2D^b^-NP396^+^ (**D**) CD8^+^ T cells isolated from the spleens of WT (n=4) and KO (n=5) mice (mean ± SD), as well as H-2D^b^-GP33^+^ (**E**) and H-2D^b^-NP396^+^ (**F**) CD8^+^ T cells isolated from the livers of WT (n=4) and KO (n=5) mice (mean ± SD). Unpaired t tests for the measurements between the two groups: *p<0.05; **p<0.01; ***p<0.001; ****p<0.0001.

**Fig. S4.**
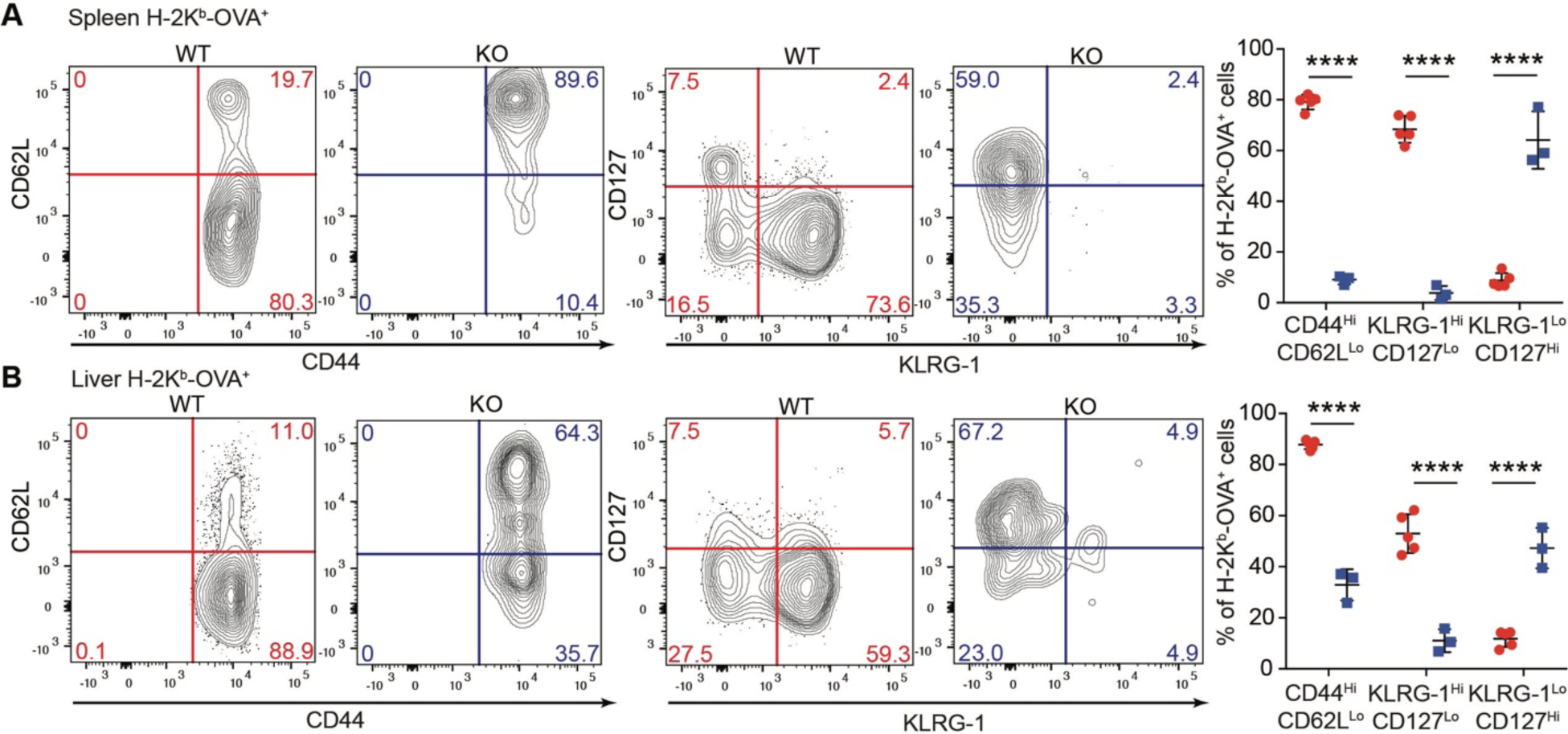
LDHA deficiency impairs CD8^+^ effector T cell response to LM-OVA infection. *Ldha*^fl/fl^ (wild-type, WT, n=5) and Tbx21^Cre^*Ldha*^fl/fl^ (knockout, KO, n=3) mice were intravenously infected with LM-OVA for 7 days. (**A** and **B**) Representative flow cytometry plots of CD44, CD62L, CD127, and KLRG-1 expression and the percentages of CD44^Hi^CD62L^Lo^, KLRG-1^Hi^CD127^Lo^, and KLRG-1^Lo^CD127^Hi^ populations among splenic (**A**) and liver (**B**) H-2K^b^-OVA^+^ CD8^+^ T cells (mean ± SD). Unpaired t tests for the measurements between the two groups: ****p<0.0001.

**Fig. S5.**
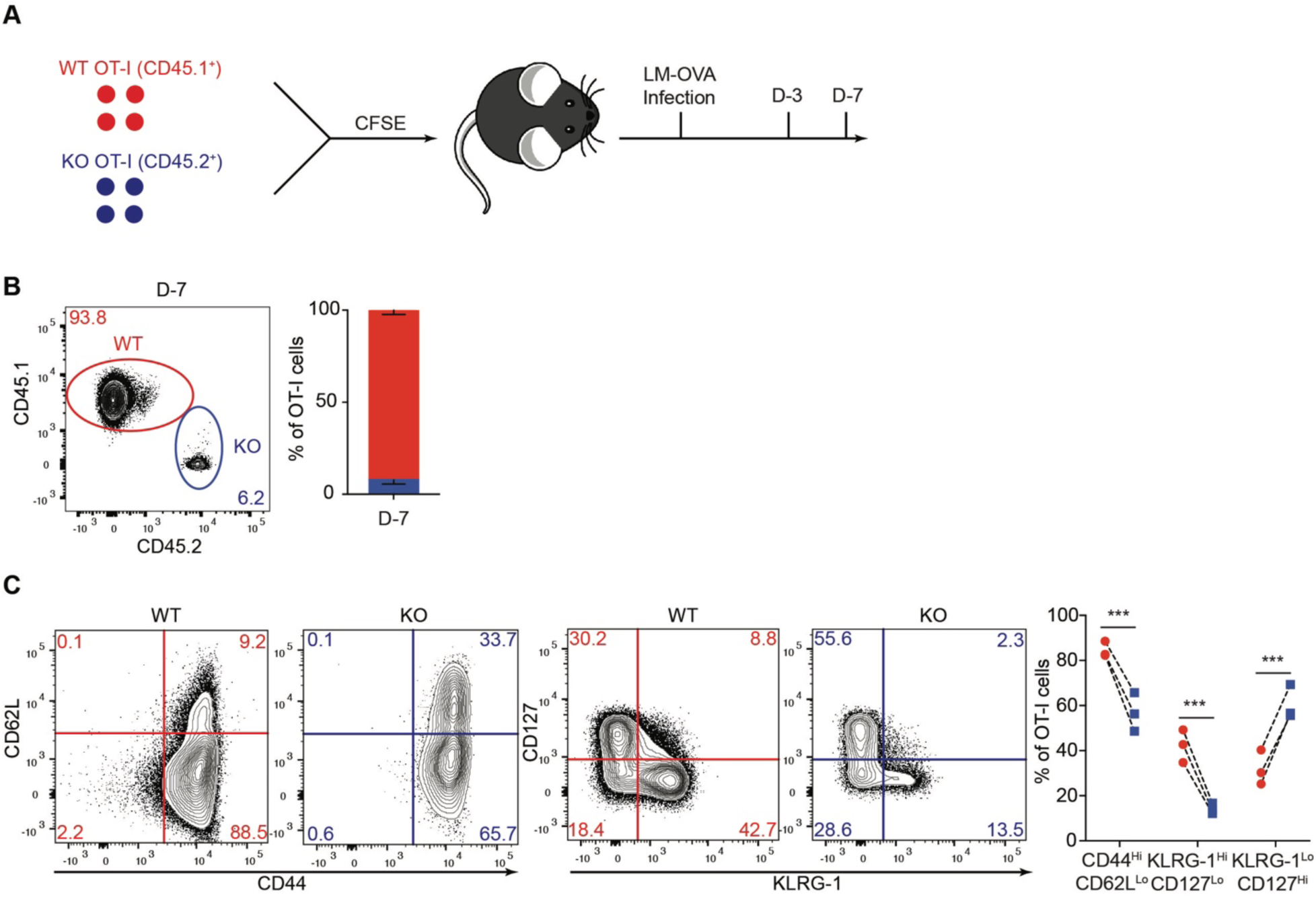
CD8^+^ effector T cell-intrinsic defects associated with LDHA deficiency. (**A**) A schematic depicting OT-I T cell co-transfer experiments. CD8^+^ T cell responses were assessed on day-3 (D-3) and day-7 (D-7) post LM-OVA infection. Briefly, congenically marked naïve OT-I T cells from *Ldha*^fl/fl^ (wild-type, WT) OT-I and Tbx21^Cre^*Ldha*^fl/fl^ (knockout, KO) OT-I mice were CFSE-labeled, mixed at a 1:1 ratio and transferred to WT recipient mice followed by infection with 5×10^3^ CFU LM-OVA. (**B**) Representative plots and ratios (mean ± SD) of liver WT and KO OT-I T cells from recipient mice day-7 post-infection (n=3). (**C**) Representative flow cytometry plots of CD44, CD62L, CD127, and KLRG-1 expression and the percentages of CD44^Hi^CD62L^Lo^, KLRG-1^Hi^CD127^Lo^, and KLRG-1^Lo^CD127^Hi^ populations among liver WT and KO OT-I T cells from recipient mice day-7 post-infection (n=3, mean ± SD). Paired t tests for the measurements between the two groups (**c**): ***p<0.001.

**Fig. S6.**
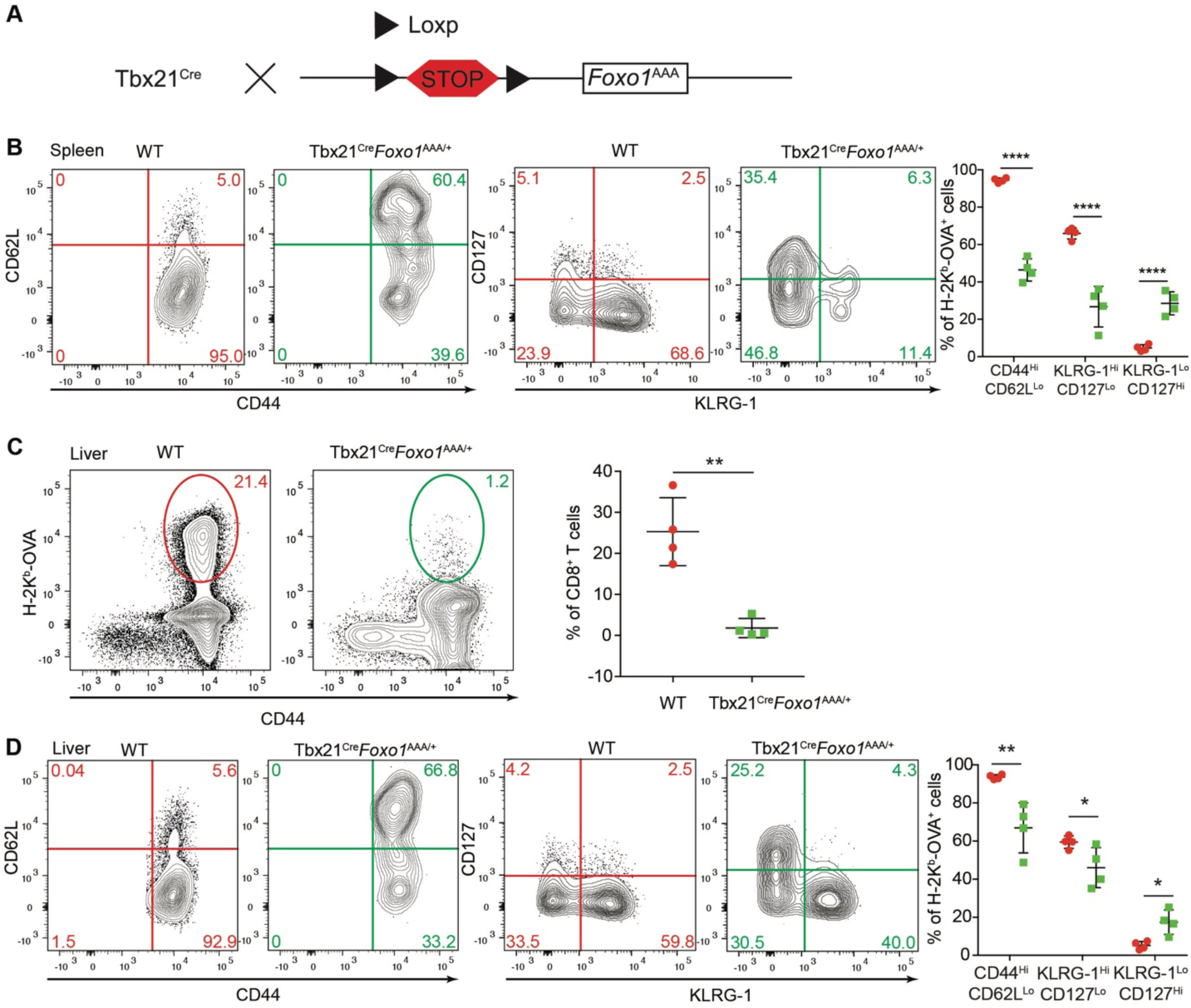
CD8^+^ effector T cell-specific expression of Foxo1^AAA^ recapitulates the T cell expansion and differentiation defects triggered by LDHA deficiency. (**A**) A schematic depicting the mouse breeding strategy and the genomic structure of *Foxo1^AAA^* mouse. (**B** to **D**) *Ldha*^fl/fl^ (wild-type, WT, n=4) and Tbx21^Cre^*Foxo1*^AAA/+^ (n=5) mice were infected with LM-OVA for 7 days. Representative flow cytometry plots of CD44, CD62L, CD127, and KLRG-1 expression and the percentages of CD44^Hi^CD62L^Lo^, KLRG-1^Hi^CD127^Lo^, and KLRG-1^Lo^CD127^Hi^ populations among splenic (**B**) and liver (**D**) H-2K^b^-OVA^+^ CD8^+^ T cells (mean ± SD). (**C**) Representative flow cytometry plots and frequencies of liver H-2K^b^-OVA^+^ CD8^+^ T cells in WT and Tbx21^Cre^*Foxo1*^AAA/+^ mice 7 days post LM-OVA infection (mean ± SD). Unpaired t tests for the measurements between the two groups (**B** to **D**): *p<0.05; **p<0.01; ****p<0.0001.

**Fig. S7.**
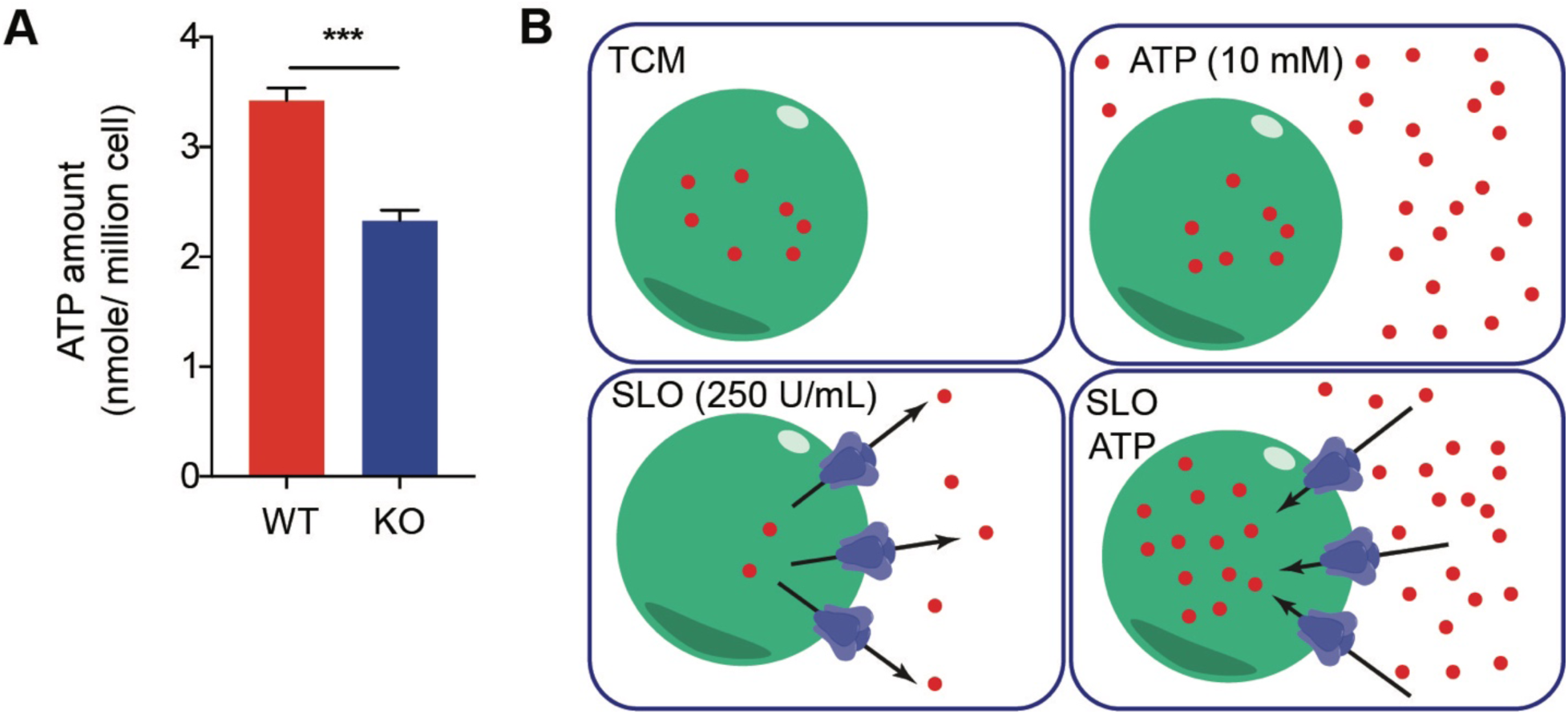
LDHA deficiency causes reduced ATP production. (**A**) Naïve CD8^+^ T cells were isolated from *Ldha*^fl/fl^ (wild-type, WT) and CD4^Cre^*Ldha*^fl/fl^ (knockout, KO) mice, and stimulated with 5 μg/ml anti-CD3, 2 μg/ml anti-CD28, and 100 U/ml IL-2 for 3 days. Cellular ATP levels were measured (n=3 per genotype, mean ± SD). (**B**) A schematic depicting a streptolysin-O (SLO)-based ATP delivery system. Unpaired t test for the measurement between the two groups (**A**): ***p<0.001.

**Fig. S8.**
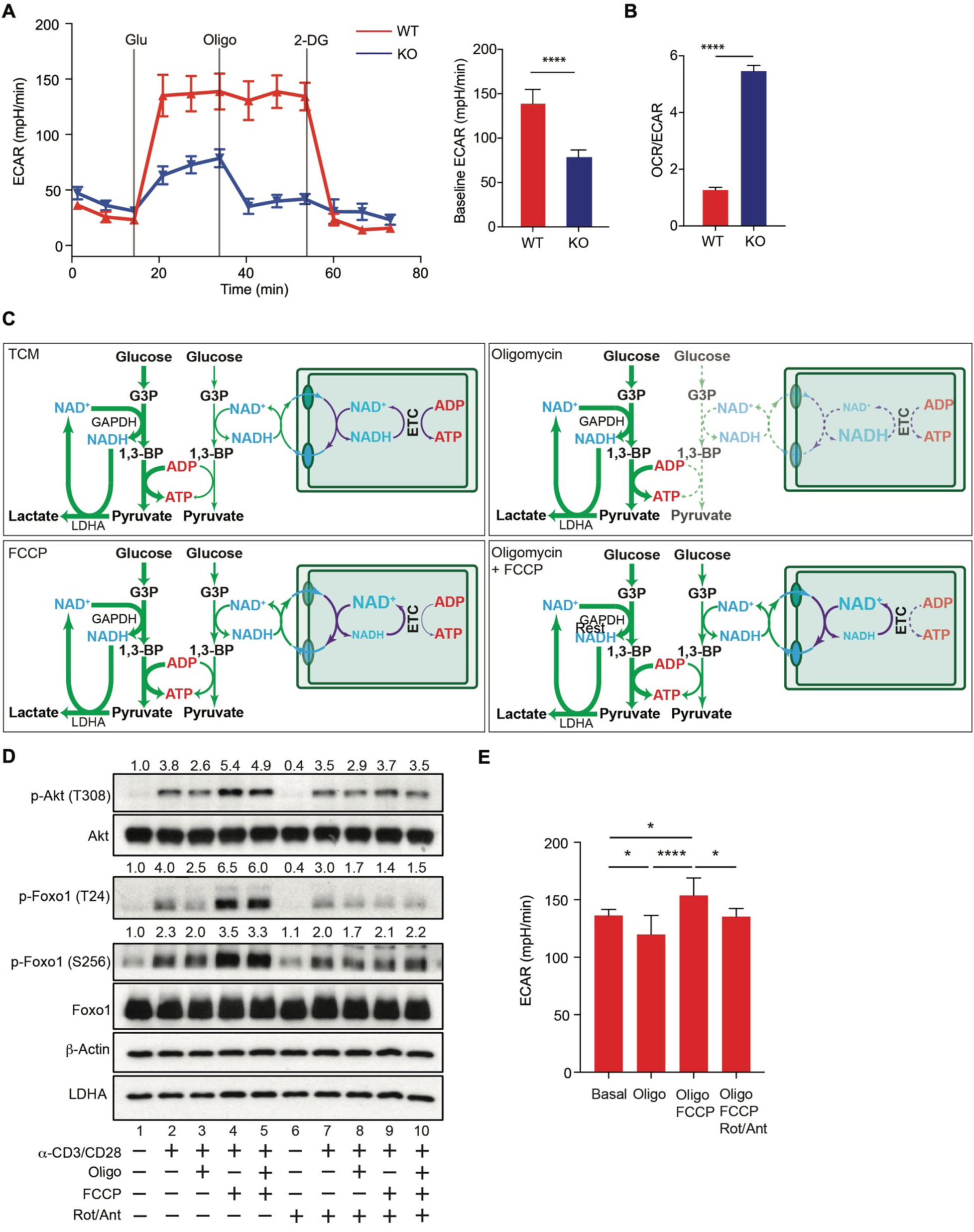
Mitochondrion electron transport chain (ETC)-mediated redox control promotes glycolysis and Akt/Foxo1 signaling. (**A**) Naïve CD8^+^ T cells were isolated from *Ldha*^fl/fl^ (wild-type, WT) and CD4^Cre^*Ldha*^fl/fl^ (knockout, KO) mice, and stimulated with 5 μg/ml anti-CD3, 2 μg/ml anti-CD28, and 100 U/ml IL-2 for 3 days before subject to measurements of extracellular acidification rate (ECAR) with a Seahorse Glycolysis stress test kit. Sequential chemical treatments are indicated as shown in the graph. Glu: glucose; Oligo: oligomycin; 2-DG: 2-deoxyglucose. (n=5 per genotype, mean ± SD). (**B**) OCR over ECAR ratios in WT and KO T cells (n=5 per genotype, mean ± SD). (**C**) A schematic of glycolysis-associated ATP generation via substrate level phosphorylation and mitochondrial ATP generation via oxidative phosphorylation as well as the redox regulatory mechanisms under the indicated treatment conditions: T cell medium (TCM), Oligomycin, FCCP, and Oligomycin + FCCP. (**D**) Day-3 activated WT T cells were collected, and incubated in RPMI medium in the absence or presence of oligomycin (Oligo) and/or FCCP and/or Rotenon (Rot) plus Antimycin (Ant). T cells were subsequently re-stimulated with biotinylated anti-CD3 and anti-CD28 through streptavidin crosslinking, and immunoblotted for p-Akt (T308), Akt, p-Foxo1 (T24), p-Foxo1 (S256), Foxo1, LDHA, and β-Actin. Normalized expression of p-Akt (T308) to Akt, p-Foxo1 (T24) or p-Foxo1 (S256) to Foxo1 were marked. (**E**) ECAR measurement for day-3 activated WT T cells treated with the indicated drugs in Seahorse Mitostress tests (n=5 per genotype, mean ± SD). Unpaired t tests for the measurements between the two groups (**A** and **B**): ****p<0.0001; two-way Anova multiple comparison tests (**E**): *p<0.05; ****p<0.0001.

**Fig. S9.**
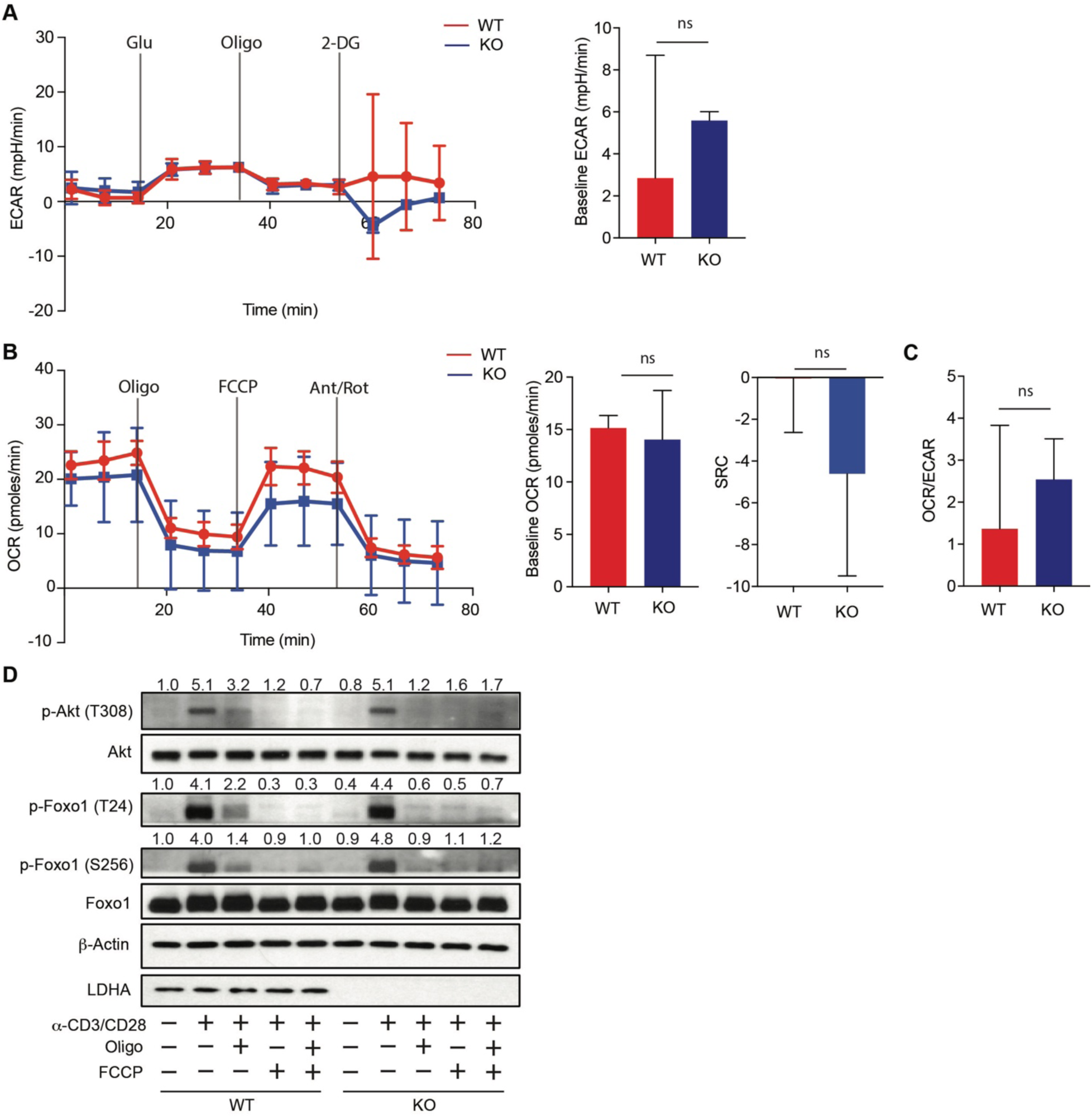
Mitochondrial ATP generation supports Akt/Foxo1 signaling in naïve T cells. (**A**) Naïve *Ldha*^fl/fl^ (wild-type, WT) and CD4^Cre^*Ldha*^fl/fl^ (knockout, KO) CD8^+^ T cells were harvested for measurements of extracellular acidification rate (ECAR) with a Seahorse Glycolysis stress test kit. Sequential chemical treatments are indicated as shown in the graph. Glu: glucose; Oligo: oligomycin; 2-DG: 2-deoxyglucose. (n=4 per genotype, mean ± SD). (**B**) Naïve WT and KO T cells were measured for oxygen consumption rate (OCR) and spared respiratory capacity (SRC) with a Seahorse Mitostress test kit. Sequential chemical treatments are indicated as shown in the graph. Oligo: oligomycin; Ant: antimycin; Rot: rotenone. (n=4 per genotype, mean ± SD). (**C**) OCR over ECAR ratios in WT and KO T cells (n=4 per genotype, mean ± SD). (**D**) Naïve WT and KO T cells were incubated in RPMI medium in the absence or presence of oligomycin and/or FCCP. T cells were subsequently re-stimulated with biotinylated anti-CD3 and anti-CD28 through streptavidin crosslinking, and immunoblotted for p-Akt (T308), Akt, p-Foxo1 (T24), p-Foxo1 (S256), Foxo1, LDHA, and b-Actin. Normalized expression of p-Akt (T308) to Akt, p-Foxo1 (T24) or p-Foxo1 (S256) to Foxo1 were marked. Unpaired t tests for the measurements between the two groups (**A** to **C**), ns: not-significant.

**Fig. S10.**
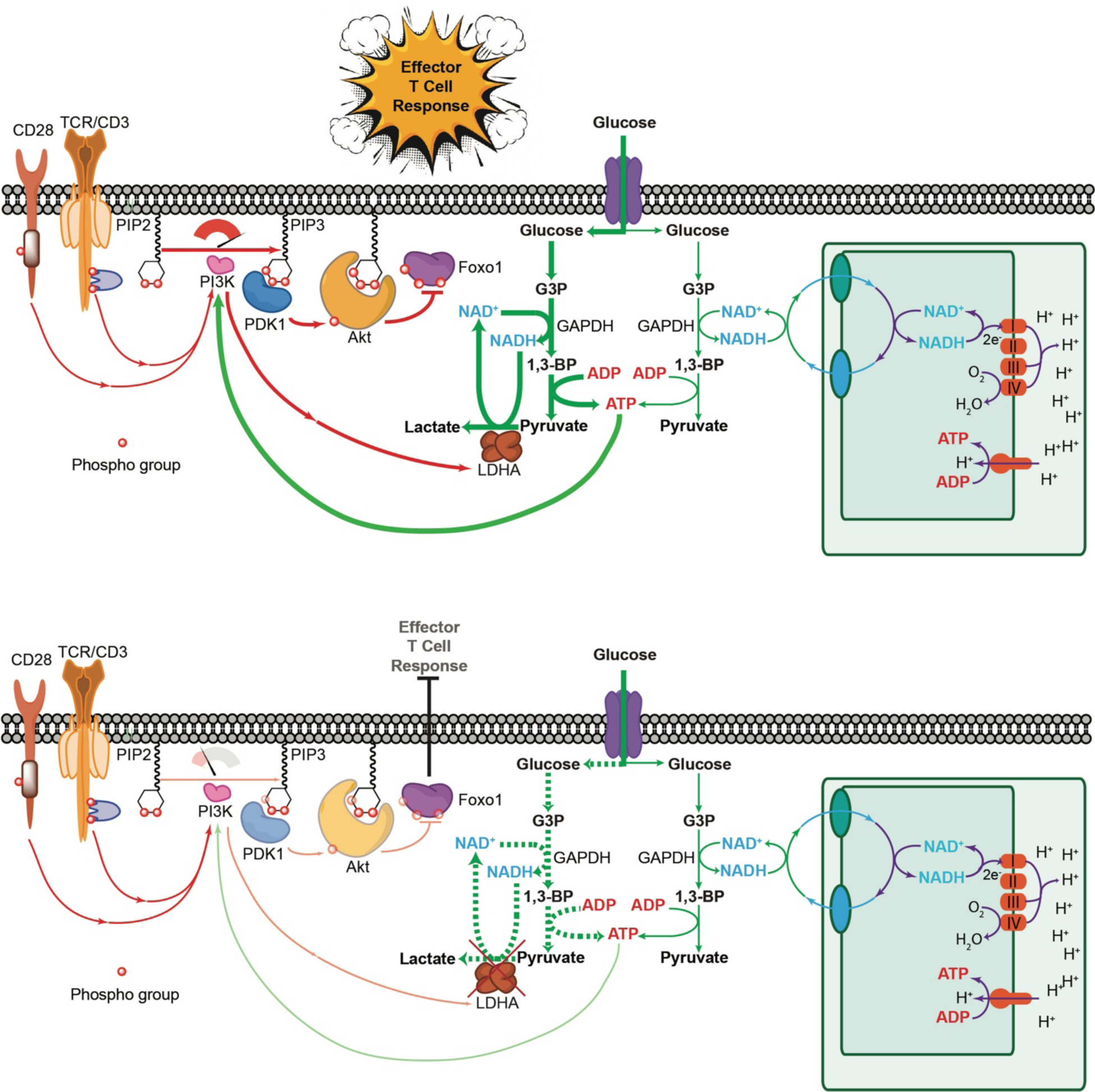
A positive feedback circuit between PI3K signaling and the LDHA-mediated glycolytic bioenergetics enables T cell immunity. Antigen receptor (TCR/CD3) and co-stimulatory receptor (e.g. CD28) signaling are integrated to activate PI3K, which promotes the expression of LDHA likely via the transcription factor c-Myc (not depicted). LDHA converts pyruvate to lactate, which regenerates NAD^+^ consumed at the GAPDH step of glycolysis, and thus maintains the redox balance and supports the high rate of ATP generation via substrate level phosphorylation. A smaller amount of cytosolic ATP can also be generated for the fraction of pyruvate not converted to lactate. In this case, NAD^+^ consumed at the GAPDH step of glycolysis is regenerated through redox shuttle systems that import electrons to mitochondria to be oxidized through the electron transport chain (ETC). ETC-mediated oxidation of NADH is associated with the generation of a proton gradient that drives the synthesis of mitochondrial ATP. Yet, cytosolic ATP, but not mitochondrial ATP, selectively fuels PI3K signaling in effector T cells. Sustained PI3K activation and PIP3 generation promotes PDK1 and Akt membrane recruitment. PDK1 exhibits constitutive kinase activity, and the PIP3-dependent recruitment of PDK1 and Akt brings them to vicinity resulting in PDK1-induced Akt phosphorylation. Activated Akt phosphorylates a number of downstream targets including the transcription factor Foxo1, repressing its activity by promoting its nuclear exclusion, which is essential for T cell expansion and effector T cell differentiation (top diagram). In the absence of LDHA, production of glycolytic ATP is severely reduced, which impairs PI3K/Akt/Foxo1 signaling causing defective T cell immunity (bottom diagram). G3P: glyceraldedyde-3-phosphate; 1,3-BP: 1,3-bisphosphoglyceric acid.

## References

1. R. Wang, D. R. Green, Metabolic checkpoints in activated T cells. Nat Immunol 13, 907–915 (2012).

2. E. L. Pearce, M. C. Poffenberger, C. H. Chang, R. G. Jones, Fueling immunity: insights into metabolism and lymphocyte function. Science 342, 1242454 (2013).

3. B. A. Olenchock, J. C. Rathmell, M. G. Vander Heiden, Biochemical underpinning of immune cell metabolic phenotypes. Immunity 46, 703–713 (2017).

4. N. M. Chapman, M. R. Boothby, H. Chi, Metabolic coordination of T cell quiescence and activation. Nature reviews. Immunology 20, 55–70 (2020).

5. M. Peng et al., Aerobic glycolysis promotes T helper 1 cell differentiation through an epigenetic mechanism. Science 354, 481–484 (2016).

6. E. H. Ma et al., Metabolic Profiling Using Stable Isotope Tracing Reveals Distinct Patterns of Glucose Utilization by Physiologically Activated CD8(+) T Cells. Immunity 51, 856–870 e855 (2019).

7. M. V. Kim, W. Ouyang, W. Liao, M. Q. Zhang, M. O. Li, The transcription factor Foxo1 controls central-memory CD8+ T cell responses to infection. Immunity 39, 286–297 (2013).

8. R. Wang et al., The transcription factor Myc controls metabolic reprogramming upon T lymphocyte activation. Immunity 35, 871–882 (2011).

9. H. L. Wieman, J. A. Wofford, J. C. Rathmell, Cytokine stimulation promotes glucose uptake via phosphatidylinositol-3 kinase/Akt regulation of Glut1 activity and trafficking. Molecular biology of the cell 18, 1437–1446 (2007).

10. N. S. Joshi et al., Inflammation directs memory precursor and short-lived effector CD8(+) T cell fates via the graded expression of T-bet transcription factor. Immunity 27, 281–295 (2007).

11. S. M. Kaech, W. Cui, Transcriptional control of effector and memory CD8+ T cell differentiation. Nature reviews. Immunology 12, 749–761 (2012).

12. J. T. Chang, E. J. Wherry, A. W. Goldrath, Molecular regulation of effector and memory T cell differentiation. Nature immunology 15, 1104–1115 (2014).

13. M. N. Navarro, D. A. Cantrell, Serine-threonine kinases in TCR signaling. Nature immunology 15, 808–814 (2014).

14. R. R. Rao, Q. Li, M. R. Gubbels Bupp, P. A. Shrikant, Transcription factor Foxo1 represses T-bet-mediated effector functions and promotes memory CD8(+) T cell differentiation. Immunity 36, 374–387 (2012).

15. R. Hess Michelini, A. L. Doedens, A. W. Goldrath, S. M. Hedrick, Differentiation of CD8 memory T cells depends on Foxo1. J Exp Med 210, 1189–1200 (2013).

16. W. Ouyang, M. O. Li, Foxo: in command of T lymphocyte homeostasis and tolerance. Trends in immunology 32, 26–33 (2011).

17. S. M. Hedrick, R. Hess Michelini, A. L. Doedens, A. W. Goldrath, E. L. Stone, FOXO transcription factors throughout T cell biology. Nature reviews. Immunology 12, 649–661 (2012).

18. W. Ouyang et al., Novel Foxo1-dependent transcriptional programs control T(reg) cell function. Nature 491, 554–559 (2012).

19. J. Rolf et al., AMPKalpha1: a glucose sensor that controls CD8 T-cell memory. European journal of immunology 43, 889–896 (2013).

20. I. Walev et al., Delivery of proteins into living cells by reversible membrane permeabilization with streptolysin-O. Proc Natl Acad Sci U S A 98, 3185–3190 (2001).

21. P. S. Costello, M. Gallagher, D. A. Cantrell, Sustained and dynamic inositol lipid metabolism inside and outside the immunological synapse. Nature immunology 3, 1082–1089 (2002).

22. C. M. Cham, T. F. Gajewski, Glucose availability regulates IFN-gamma production and p70S6 kinase activation in CD8+ effector T cells. J Immunol 174, 4670–4677 (2005).

23. P. M. Gubser et al., Rapid effector function of memory CD8+ T cells requires an immediate-early glycolytic switch. Nature immunology 14, 1064–1072 (2013).

24. P. C. Ho et al., Phosphoenolpyruvate Is a Metabolic Checkpoint of Anti-tumor T Cell Responses. Cell 162, 1217–1228 (2015).

25. A. V. Menk et al., Early TCR Signaling Induces Rapid Aerobic Glycolysis Enabling Distinct Acute T Cell Effector Functions. Cell reports 22, 1509–1521 (2018).

26. M. D. Buck et al., Mitochondrial Dynamics Controls T Cell Fate through Metabolic Programming. Cell 166, 63–76 (2016).

27. Z. A. Knight, K. M. Shokat, Features of selective kinase inhibitors. Chemistry & biology 12, 621–637 (2005).

28. J. R. Somoza et al., Structural, biochemical, and biophysical characterization of idelalisib binding to phosphoinositide 3-kinase delta. The Journal of biological chemistry 290, 8439–8446 (2015).

29. M. D. Goncalves, B. D. Hopkins, L. C. Cantley, Phosphatidylinositol 3-Kinase, Growth Disorders, and Cancer. The New England journal of medicine 379, 2052–2062 (2018).

30. O. Warburg, The Metabolism of Carcinoma Cells. The Journal of Cancer Research 9, 148–163 (1925).

## References

1. M. Peng et al., Aerobic glycolysis promotes T helper 1 cell differentiation through an epigenetic mechanism. Science 354, 481–484 (2016).

2. W. Ouyang et al., Novel Foxo1-dependent transcriptional programs control T(reg) cell function. Nature 491, 554–559 (2012).

3. A. G. Levine et al., Stability and function of regulatory T cells expressing the transcription factor T-bet. Nature 546, 421–425 (2017).

4. I. Walev et al., Delivery of proteins into living cells by reversible membrane permeabilization with streptolysin-O. Proc Natl Acad Sci U S A 98, 3185–3190 (2001).

